# A potential acoustic role for CFTR ion channel in conductive hearing loss

**DOI:** 10.1101/2023.09.23.559053

**Authors:** Pramodha Liyanage, Kyu-Shik Mun, Gianni Carraro, Herbert Luke Ogden, Yunjie Huang, Jesun Lee, Yashaswini Ramananda, Barry R Stripp, Kavisha Arora, Nathan Salomonis, Lisa L. Hunter, Anjaparavanda P. Naren

## Abstract

Loss-of-function mutations in the cystic fibrosis transmembrane conductance regulator (CFTR) gene cause cystic fibrosis (CF). The middle ear and eustachian tube could be adversely affected in CF. In this study, we provide evidence of the role of CFTR function in conductive hearing. We developed an in-situ model to determine CFTR dependent fluid secretion in the middle ear using native mouse auditory capsule. A unique middle ear-on-a-chip was developed to address the functional and molecular basis of conductive hearing impairment. Using single-cell transcriptomics, middle ear cell composition and the associated transcriptomic signature were compared between CF and WT groups. A specialized subset of epithelial cells expressed CFTR with an overlapping signature with secretory epithelial cells. Genes related to ciliogenesis, hearing and ossification were significantly altered in CF mice middle ear. Our data suggest that CF middle ear may be at higher risk for conductive hearing loss.

## Introduction

Auditory perception plays a crucial role in communication, learning, integration, survival, and evolution. Hearing is one of the most developed senses in higher organisms and is specifically dependent upon the accurate function and synchronization of the auditory system. The ear drum captures sound energy and transmits it to the sensory hair cells in the inner ear through the air-filled middle ear cavity. The three small bones (ossicles), malleus, incus and stapes in the middle ear, form a semi-rigid chain of bones that transmit the sound waves as a form of vibration from tympanic membrane (TM) to the inner ear^1^. The middle ear cavity connects to the nasopharynx through a small canal called the Eustachian tube (ET) which ventilates and drains the middle ear cavity^2^. The ET regulates pressure between the middle ear and pharynx, which is essential for normal hearing. The most common cause of conductive hearing loss is otitis media with effusion (OME), in which mucus and fluid buildup in the middle ear^3^. Other conditions may also adversely affect middle ear conductive hearing. For example, dysfunction of the eustachian tube, blunt or penetrating trauma to the ear, ossicular erosion, cholesteatoma and otosclerosis may contribute to middle ear conductive hearing loss (CHL) ^4^.

The middle ear epithelium is primarily responsible for regulating the middle ear mucosa. The natural mechanisms for maintaining a healthy tympanic cavity include mucociliary clearance, removal of middle ear secretions by pumping action of the ET, and transepithelial fluid transport^5–8^. Ion channels also play a particularly important role in middle ear function by regulating the volume and composition of the periciliary fluid in the middle ear cavity^9,10^. The Cystic Fibrosis Transmembrane Conductance Regulator (CFTR), an anion channel, is one of the most important channel proteins in the secretory epithelia; it plays a critical role in regulating anion and fluid transport across the apical membrane^11,12^. Mutations of the *CFTR* gene lead to defective CFTR protein and abnormal CFTR function causing cystic fibrosis (CF). Specifically, Cl^-^ ion secretion is reduced, and sodium ion absorption is activated. Consequently, individuals with CF have epithelial secretions that become sticky and thick^13^. More than 2000 CFTR mutations have been discovered which are classified typically into six different groups. These mutations either affect CFTR folding, stability or trafficking to the plasma membrane, reducing the number of functional CFTR channels on the cell membrane^14^. The complexity of CFTR mutations has complicated the search for a universal drug to treat CF. The most common CF mutation, ΔF508, disrupts CFTR folding and is found in ∼87% CF patients. CF patients are highly vulnerable to recurrent pulmonary infections by gram negative bacteria, mainly *Pseudomonas aeruginosa,* because poor mucociliary clearance allows bacteria to more readily colonize the airway.

While almost all CF patients develop chronic respiratory tract infections, most are believed to develop middle ear infections as well. However, the prevalence of middle ear infections among CF patients remains unclear^15–17^. Some of the previous studies have shown that a lower rate of inflammatory middle ear disease in CF patients compared to controls ^16^, while others have suggested that CF patients are not at higher risk than controls ^17^. However, most studies were not specifically designed to test for middle ear disorders, such as middle ear infection or CHL. Several recent studies conducted with traditional audiometric and otologic techniques found that the CF group had a significantly higher occurrence rate of OME and CHL compared to the control group^18,19^. In CHL due to OME, sound energy conduction may be attenuated by as much as 30-40 dB^20^. Currently, the exact causes and mechanisms of CHL in CF patients are not well understood.

In this study, we examine the expression and activity of CFTR in the middle ear and the impact of the most common CF mutation, ΔF508, on middle ear conductive hearing loss. To study the expression of functional CFTR in the middle ear, a novel middle ear *in-situ* CFTR functional characterization method was developed using the entire auditory capsule. This study was advanced in two different ways to evaluate the effect of CFTR on sound energy propagation in the middle ear. First, a middle ear model was developed including a TM, a middle ear cavity with epithelial cells, and an ET; this model allowed us to quantify the effect of CFTR-expression in middle ear epithelia, and better understand sound energy absorption in relation to CFTR expression. Second, single cell RNA sequencing (scRNA-seq) technology was used to compare normal and ΔF508 mutant CFTR middle ear transcriptome to identify possible genomic level changes that may lead to CHL.

## Results

### CFTR function in the middle ear augments mucociliary clearance

To demonstrate the functional signature of CFTR in the middle ear, an *in-situ,* physiologically relevant model was created using native mouse middle ear tissue. First whole auditory capsule containing mouse middle ear and ET was micro-surgically isolated (Fig. 1a, a cross section of ΔF508 mouse middle ear is shown in Extended Data Fig. 1). One end of the auditory capsule contained the outer ear opening and the other end had the ET opening. Peripheral tissues were carefully removed to clear bone from each end of the preparation. After meticulously removing the TM and ossicles by microdissection, the middle ear cavity and the ET made a continuous tube-like structure with two openings. The rationale for this procedure is to facilitate a flow of buffer (using a peristaltic pump) from the outer ear opening to the ET through the middle ear cavity (Fig.1b). The outer ear opening of the auditory bulla was connected to a tube to perfuse the buffer through the middle ear cavity (Fig. 1b and 1c). The detailed procedure is described in the methods section. Using this design, we perfused the buffer through the middle ear cavity and collected it from the ET (Fig. 1d).

**Fig. 1.**
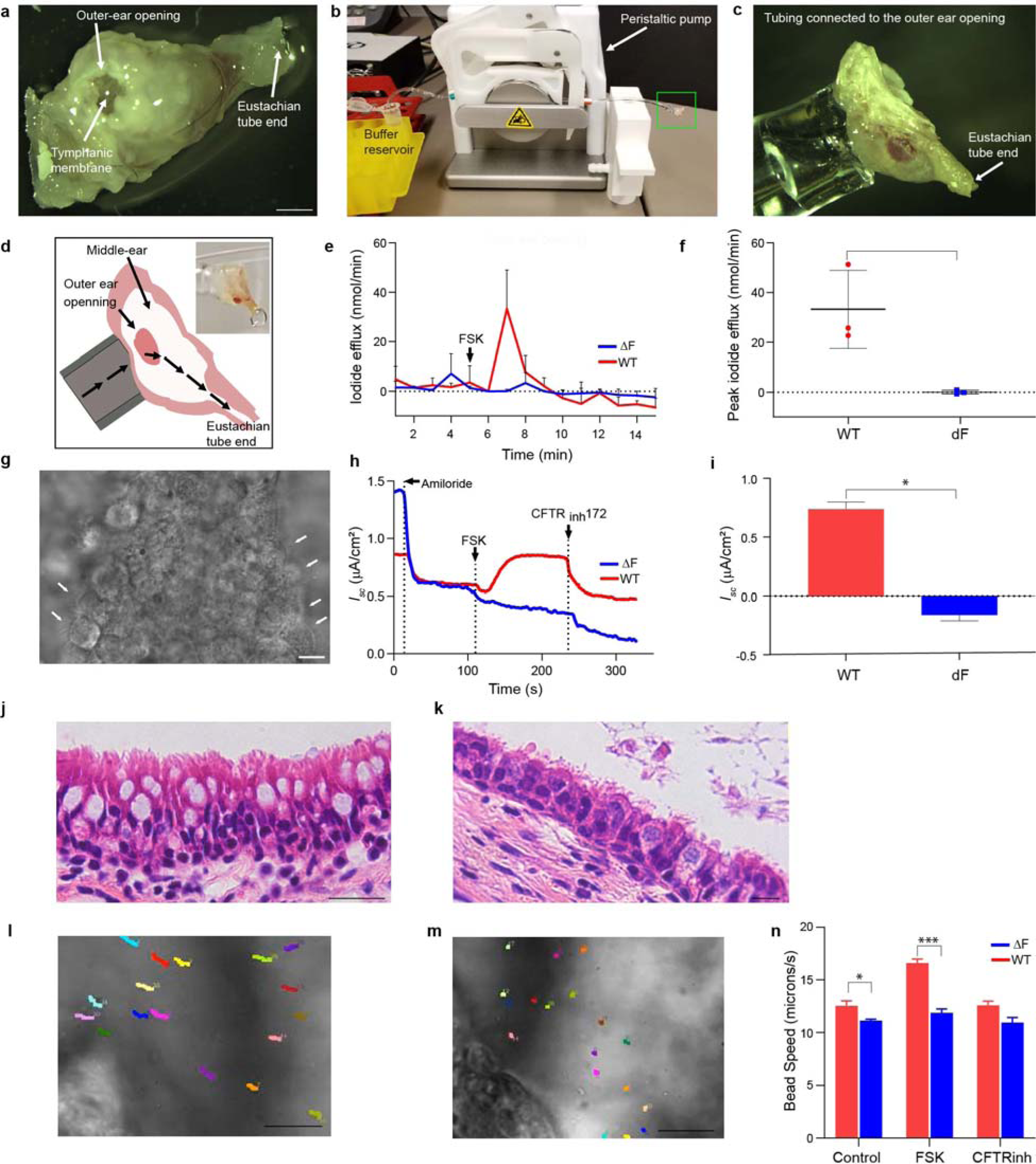
CFTR function in the middle ear facilitates mucociliary clearance. **a,** Morphology of the isolated mouse auditory capsule. Scale bar is 1 mm **b,** The experimental setup for the middle ear perfusion system. The tubing connected to the peristaltic pump drives the buffer from the receiver to middle ear. The green box is enlarged in c. **c,** The outer ear opening is attached to the tubing. **d,** A schematic diagram depicting the buffer flow through the middle ear cavity and ET. The buffer enters to the middle ear through the outer ear opening. After completely filling the whole middle ear, buffer moves out from the ET-end opening. **e,** Iodide efflux measurement of whole middle ear and ET. Both WT and ΔF508 mice middle ear/ET was tested for forskolin induced CFTR activity. **f,** Comparison of the peak Iodide efflux response between WT and ΔF508 mutant mice middle ear/ET. **g,** A phase contrast image of fully differentiated mouse middle ear/ET primary cells on transwell inserts. The white arrows indicate the ciliated cells. Scale bar is 50 μm. **h,** CFTR-mediated short-circuit current (*I_sc_*) in middle ear/ET monolayers in Ussing chamber. Representative traces show the sequential addition of Amiloride, forskolin (FSK) and CFTR inhibitor 172. **i,** Comparison of the forskolin induced peak CFTR response for short-circuit current (*I_sc_*) experiment in middle ear/ET. **j,** A cross section of the WT mouse middle ear showing the ciliated, pseudostratified epithelia. Scale bar is 20 μm. **k,** The morphology of the ΔF508 mutant mouse middle ear epithelia. Scale bar is 20 μm. **l,** Cilia generated, directional bead flow of WT middle ear cells. Scale bar is 5 μm. **m,** The ΔF508 mutant middle ear shows an impaired bead flow. Scale bar is 5 μm. **n,** Quantitative comparison of the cilia-generated bead flow between WT and ΔF508 mutant mice. The bead flow was enhanced in response to FSK only in the WT middle-ear.

Flux studies with ion selective electrodes (referred as iodide efflux assay) are a well-established method to evaluate CFTR function^21^. The iodide efflux assay was performed with the novel experimental setup to evaluate CFTR activity in the middle ear/ET (Fig. 1e). The WT mouse middle ear/ET showed a clear CFTR response peaking at the 7^th^ minute after addition of FSK, a cAMP agonist (Fig. 1e red line). Given that the ΔF508 CFTR mutant does not express functional CFTR at the plasma membrane, a similar iodide efflux assay was performed in ΔF508 mouse middle ear/ET as a negative control. The blue line in Figure 3e shows the time course of ΔF508 mutant mouse middle ear/ET; as expected, FSK stimulation was not associated with CFTR activity. it was concluded that WT mice have increased CFTR activity, indicating the presence of active CFTR in the middle ear/ET (Fig. 1f).

Mouse middle ear/ET epithelial cells were isolated and cultured on transwell membranes using an air-liquid interface in which epithelial cells represent morphological characteristics akin to the cells in the native mouse middle ear epithelium (Fig. 1g). Middle ear epithelial cell isolation and culturing protocols have been described previously ^22–24^. To further validate the presence and function of CFTR in the middle ear/ET, middle ear primary cells were cultured and tested for CFTR-dependent short circuit current (*I_sc_*) using an Ussing chamber (Fig. 1h). CFTR-dependent *I_sc_* traces were obtained for WT middle ear epithelial cells in response to amiloride (sodium channel inhibitor), forskolin and CFTR inhibiter (CFTR_inh172_); ΔF508 cells did not respond to forskolin (Fig 1i).

Next, cilia-driven, periciliary fluid flow which is important for mucus clearance in the middle ear/ET was investigated. Defects in the periciliary fluid flow lead to severe disease manifestations due to the accumulation of bacteria/viruses and particulate matter in the middle ear. Middle ear cilia distribution, and coordinated polarity toward the ET, have been reported^25^. However, the role of CFTR in cilia-driven middle ear fluid flow has not been studied. Figure 1j and 1k illustrate cilia distribution in the middle ear epithelium of WT and ΔF508 samples respectively. The cilia-driven fluid flow was quantified and compared using fluorescence beads (Fig. 1l, 1m) to study the effect CFTR on ciliary beat dynamics. Bead flow was monitored on the native middle ear epithelium (Supplementary video 1 and 2). Under the control condition (i.e., unstimulated state), bead speed was significantly higher (p<0.05) in WT than ΔF508 preparations. When CFTR was stimulated using FSK, bead speed increased rapidly (p<0.001), indicating that cilia-generated flow is directly related to CFTR activation (Fig 1l). When CFTR activity was inhibited, cilia-generated bead flow returned to normal levels.

### CFTR localizes to the apical PM of polarized middle ear epithelial cells

The middle ear cavity and ET are covered by epithelial cells that are contiguous with the upper airway. The middle ear epithelium is composed of basal, secretory, non-secretory, and ciliated cells. The ET is lined by a pseudostratified epithelium that extends from the opening of the ET towards ventral part of the middle ear (hypotympanum) (Fig 2a). A simple epithelium is found in the dorsal part (epitympanic region) of the middle ear cavity^26–28^. The middle ear/ET mucosa provides protection to the middle ear; this epithelium is covered by a thin film of periciliary fluid^6^. Ion and water homeostasis in the middle ear epithelia is critical in modulating periciliary fluid volume and maintaining a fluid-free middle ear cavity. CFTR plays a major role in hydration of periciliary fluid, maintenance of pH, and regulation of other channels and immune responses in the respiratory track epithelia^29^. To investigate the expression of CFTR in the middle ear, RNA *in-situ* hybridization method was used (RNAscope, Advanced Cell Diagnostics), with a unique CFTR probe set (see methods). It was demonstrated that CFTR puncta was significantly higher in WT middle ear/ET epithelium (Fig 2b) than in a CFTR-deficient state. Furthermore, an RNAscope probe was used to detect MUC5b mRNA as a positive control, which is abundant in the middle ear epithelial cells (Fig 2b). In parallel, the CFTR expression pattern was determined using the peptide affinity purified mouse anti-CFTR 3194 antibody (obtained from Dr. Marino, Memphis, TN, which has been extensively validated) ^30^. Immunofluorescence analysis indicates that wildtype CFTR protein is localized to the apical membrane of epithelial cells residing in the middle ear/ET region (Figure 2c). However, CFTR distribution was limited to the ventral side of the middle ear cavity. Interestingly, no CFTR protein was detected within middle ear/ET epithelium of ΔF508 preparations (Fig. 2c lower panel).

**Fig. 2.**
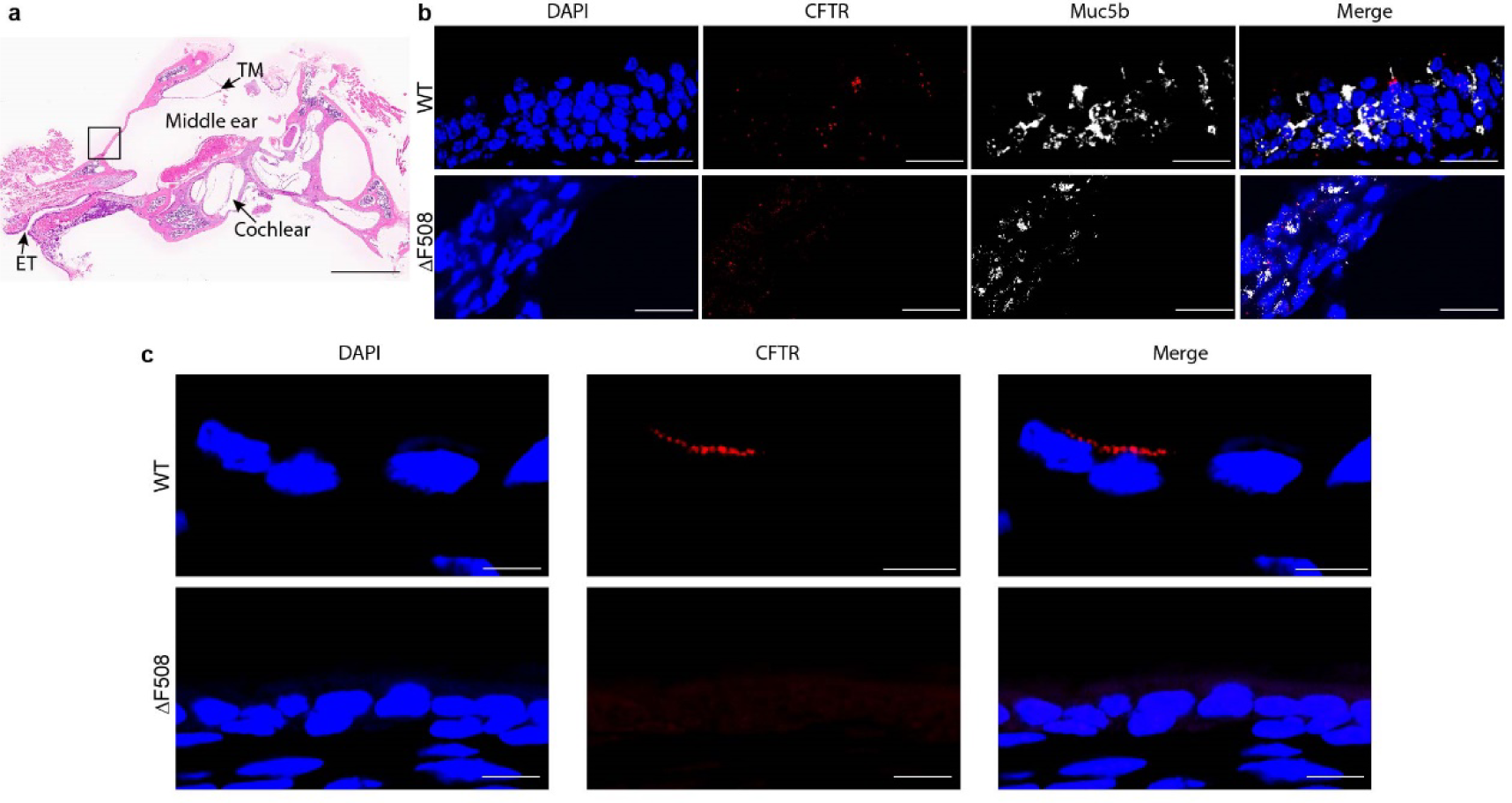
Immunofluorescence staining confirms CFTR expression in middle ear epithelium. **a,** Histology showing anatomical features of WT mouse middle ear. TM-Tymphanic membrane, ET-Eustachian Tube. Scale bar is 1 mm. **b,** CFTR mRNA detection by RNA-scope method. Both WT (top) and ΔF508 (bottom) middle-ear sections were hybridized with probes set to detect CFTR (red) Muc5b (white). The nuclei were stained with DAPI. scale bar is 25 μm. **c,** CFTR detected in the WT mouse middle-ear (top panel) by anti-CFTR antibody (red) in the area marked (black box) in the “a”. Scale bar is 5 μm.

### Designing and fabrication of middle ear on-a-chip to demonstrate a role of CFTR in sound propagation

The pathophysiological effects of loss of CFTR expression and function have been widely studied in the context of lungs, gut, pancreas, liver and kidney^31–33^, with very few reports citing the role of CFTR function in the middle ear^6^. We rationalized that CFTR controls cilia-generated flow, and transepithelial fluid transport, to maintain the physiological level of periciliary fluid on middle ear/ET mucosa. Mutant CFTR is thought to cause dehydration of the air liquid interface (ALI) and thickening of the mucus layer, altering the volume and composition of the periciliary fluid lining the middle ear/ET mucosa. We further hypothesize that loss of CFTR function leads to abnormal ALI on the middle ear/ET epithelial lining and perturbs middle ear acoustic functions.

To determine if epithelial pathology in CF affects sound energy propagation in the middle ear, a middle ear/ET model was developed using 3D printing technology (Fig 3a). Only the TM, middle ear and ET were included in the model to exclusively evaluate the influence of middle ear/ET epithelia on middle ear acoustics. The middle ear/ET model can be segmented into two parts. First, fully differentiated, middle ear epithelial cells are grown on an ALI-containing transwell insert which resembles the tympanic cavity. The same set of transwell inserts were used for measuring CFTR-dependent short circuit currents in Figure 1g and 1h. In the second piece, the outer ear and ET are modeled to a single unit that was then inserted into the set of transwell inserts. This part was casted as a single piece guided by a 3D-printed model (see methods for details, Extended Data Fig. 2). The outer ear section was a hollow, cylindrical tunnel with an inner diameter of 10 mm; one end is open to air and the other end is connected to the TM. The TM was made by very thin layer (5 μM) of PDMS and attached to the inner end of the outer ear canal separately (Fig. 3b). To measure middle ear sound conduction, the ear tip of a tympanometer probe was inserted into the outer ear (Fig. 3a). The final version was developed through multiple iterations and had excellent biomechanical properties that were similar to the intact human middle ear. Wideband energy absorbance was used to characterize the effect of middle ear epithelial properties and to evaluate the sensitivity of the epithelium model. This technique, referred to as Wide Band Tympanometry (WBT), uses electroacoustic immittance to measure middle ear function across a physiologically relevant frequency range for normal human hearing sensitivity. It measures the acoustic energy absorbed by the middle ear with respect to the incident energy, as a function of pressure induced in the outer ear canal. WBT can provide valuable information about conduction of sound energy through the ear, and is sufficiently sensitive to distinguish different middle ear pathologies such as ossicular fixation and disarticulation, mastoid disease and tympanic membrane perforation^34,35^. The model was tested in three forms: non-cell (collagen coating only), non-ciliated cells and ciliated cells (Fig. 3c). The absorbance function between 250-8000 Hz was recorded at ambient and peak pressure three times in each condition to assess reliability. The three experimental conditions showed significantly different tympanogram patterns at high frequencies at ambient pressure (5000 Hz-7000 Hz). The collagen coated membrane (control) showed low absorbance at high frequencies, while absorbance was improved with non-ciliated and ciliated cells in the same frequency range. The ΔF508 mutant epithelial cells showed a significant drop in absorbance at 3000 Hz and continued to have lower absorbance at higher frequencies compared to WT. Next, the model with collagen coating was tested in three conditions to mimic serous OME (Extended data Fig. 3). The baseline condition was measured, then the middle ear component of the model was partially filled with distilled water and re-measured. Last, the model’s middle ear cavity was completely filled with distilled water and absorbance re-measured. In the partial condition, a mass-loading effect was observed, with large decrease in the high frequencies, and resonant peaks at 800 and 1800 Hz. In the completely filled condition, absorbance was very low across all frequencies, due to combined stiffness and mass-loading effects. This experiment demonstrated similar effects for WBT measurements as seen in humans with partial serous OME and with the middle ear space completely filled with OME. These data confirmed that the model is sensitive to the physiological changes of the epithelium that help to distinguish different pathological conditions. However, further evaluation is needed to understand tympanogram changes based on the epithelial condition and how it relates to CHL.

**Fig. 3.**
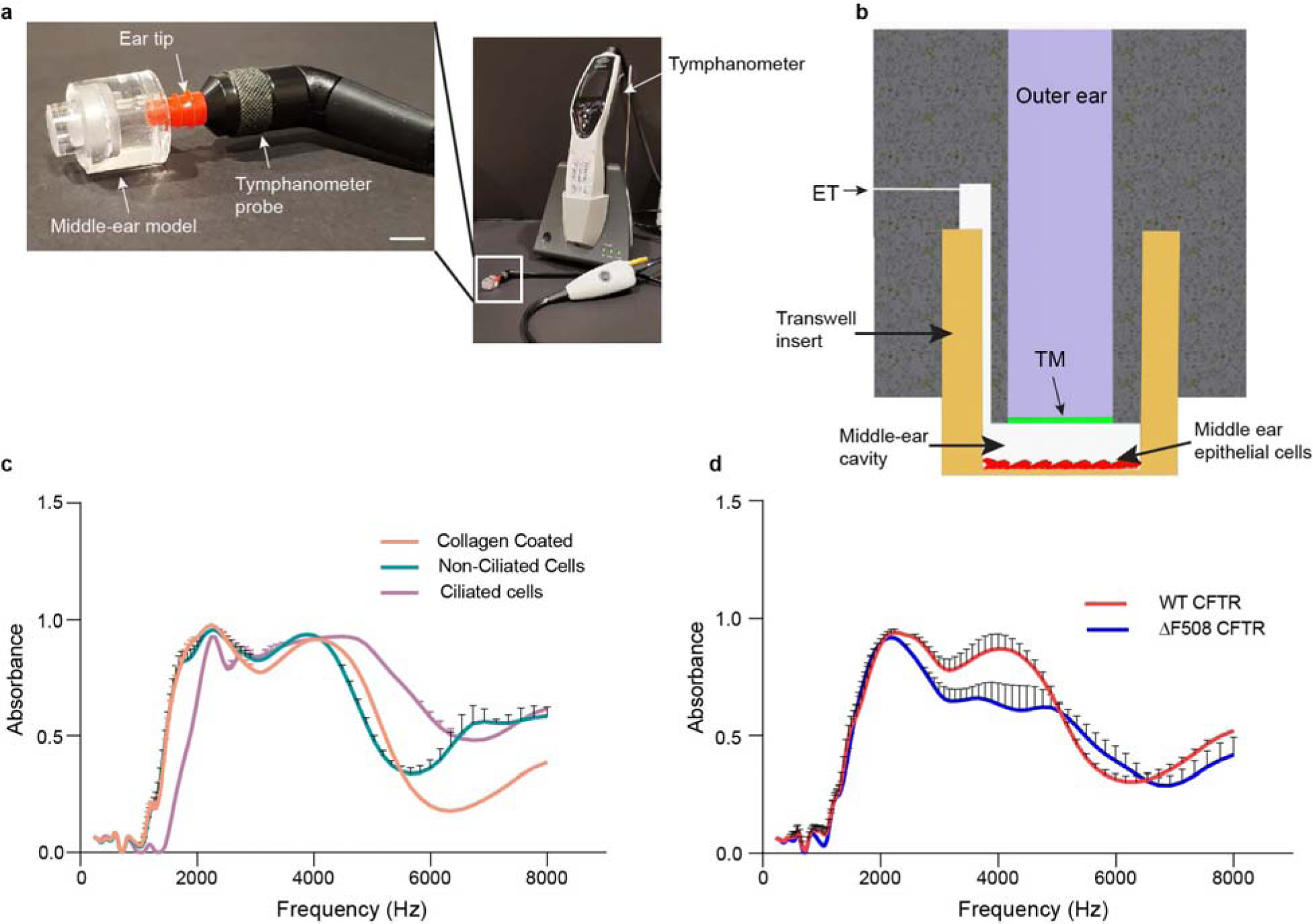
Custom made microfabricated middle ear-on-a-chip reveals perturbed middle ear absorbance in CF middle ear epithelium with implications to conductive hearing loss. **a,** The middle ear chip connected to the tympanometer. The ear tip (red) is inserted into the outer ear cannel. Scale bar is 5 mm. **b,** Schematic representation of a cross section of the middle ear design (ET-Eustachian tube, TM-tympanic membrane). The semi-permeable transwell with cultured middle-ear epithelial cells is inserted into the 3D printed model. **c,** Tympanometry measurements were sensitive to distinguish different conditions (Collagen coated: No cells; non-ciliated cells: planar non-polarized Human Embryonic Kidney Cells; Ciliated cells: Differentiated and polarized ciliary sub-type enriched airway epithelial cells). **d,** Middle ear absorbance was measured using middle-ear-on-a-chip with cultured WT and ΔF508 *<u>cftr</u>* middle ear epithelial cells.

### Single cell RNA sequencing uncovers cellular and genetic heterogeneity leading to conductive hearing loss

Although the physiological and electrophysiological results are critical in evaluating CFTR function in the middle ear, these data do not provide information on specific cell populations and gene expression that can be scaled for disease modeling. To assess the molecular and cellular impacts of ΔF508 mutant CFTR on the predominant middle ear/ET cell populations, scRNA-Seq was performed on total cells isolated from the middle ear of either ΔF508 or WT animals (Table S1). Unsupervised analysis of ∼15,000 combined cells (using Iterative Clustering and Guide-gene Selection (ICGS2) software) identified 35 cell populations, each defined by unique marker genes (Fig. 4a and Table S3-S4). Hematopoietic cells were the dominant population, with little difference in the overall distribution in both genotypes (Extended Data Fig. 4a).

**Fig. 4.**
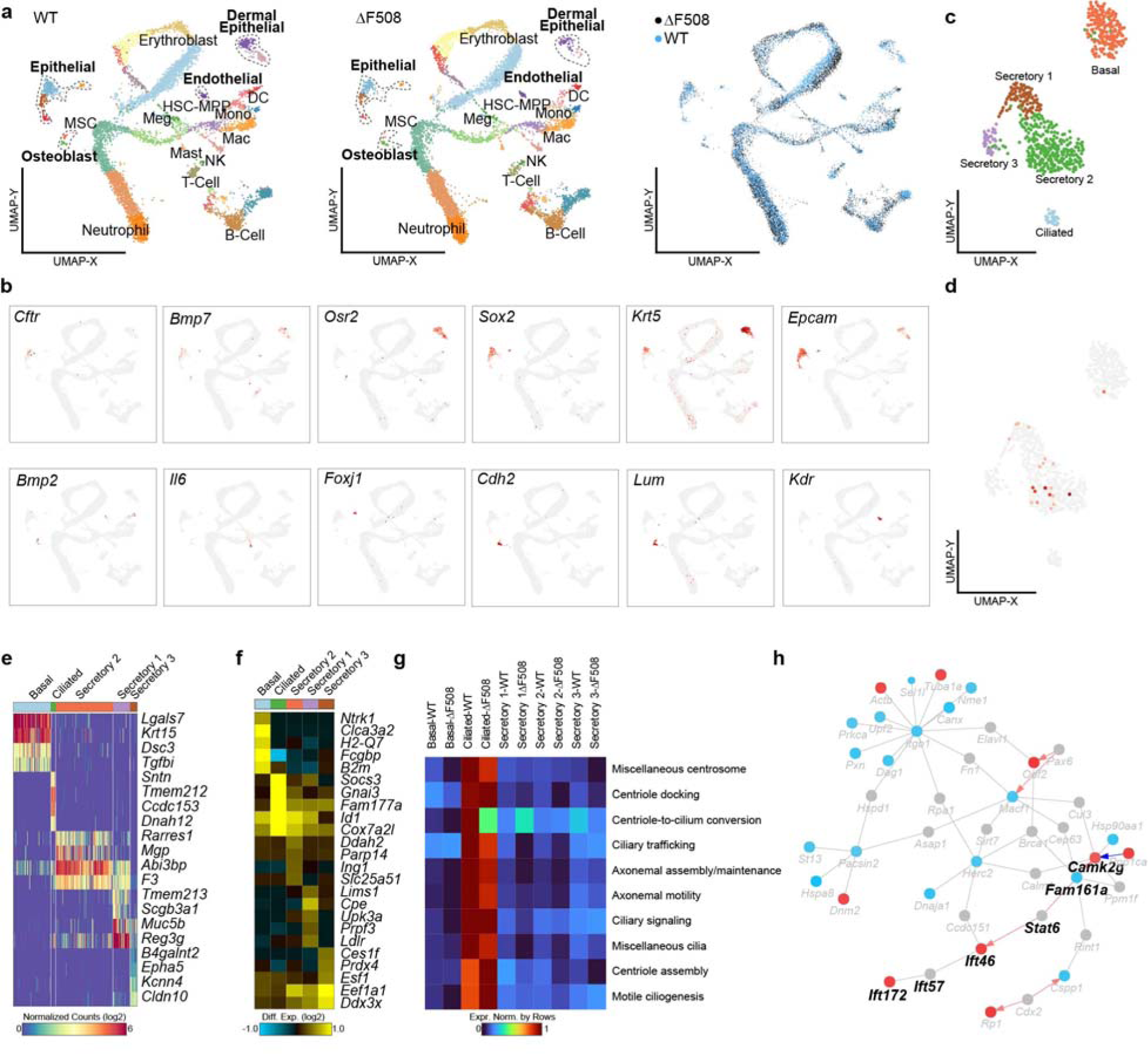
Single-cell RNA sequence analysis reveals extensive transcriptional alterations in ΔF508 *Cftr* middle ear. **a,** UMAP projection of middle ear/ET single-cell populations defined by the software ICGS2 following joint-analysis of control and ΔF508 CFTR mutant captures. Separate plots for control (left), ΔF508 (middle) and combined (right) cells. **b,** UMAP projection of gene expression for selected epithelial, endothelial and osteoblast markers genes (grey = no expression, red = high). **c,** UMAP of EPCAM^+^ epithelial cell sub-clusters. **d,** CFTR expression is limited to secretary 1 and 2 subpopulations. **e,** Heatmap displaying epithelial cluster specific markers. **f,** The top significantly differentially regulated genes in the ΔF508 compared to WT cells. **g,** Heatmap of average gene expression in WT and ΔF508 epithelial cells for ciliogenesis Gene Ontology terms. The most cilia related gene expression is limited to the ciliated cell sub-cluster and are decreased in the ΔF508. **h,** Predicted gene interactions in cilia, comparing ΔF508 to WT. DEGs were restricted to those in cilia related Gene Ontology terms with indirect interactions. The red, blue and grey dots represent the downregulated, upregulated and unaltered genes respectively (empirical Bayes t-test p<0.05). Red arrows denoted predicted transcriptional regulation (PAZAR/TRRUST).

Initially broad cell population labels from gene-set enrichment were confirmed based on the expression of well-defined marker genes for epithelial, osteoblast and endothelial cells (Fig. 4b). Broad differences in gene expression were found when comparing the ΔF508 to equivalent matching WT cells for all cell population pairs. Specifically, there was broad upregulation of genes that induce apoptosis (e.g., *Bnip3l*), upregulation of cell-cycle promoting genes in erythroblast and monocyte progenitors, and extensive upregulation of genes in B-cell and neutrophil cell populations, and down-regulation of genes involved in oxidative damage *(e.g., Gpx1)* (Extended Data Fig. 4b and Table S5-S7). These noted cell populations were associated with perturbation of core lineage specification (B-cell), cell stress and hypoxia (neutrophil), and epigenetic/polycomb (erythroblast) associated-predicted transcriptional regulatory networks (Extended Data Fig. 4c). Conversely, when we examined non-hematopoietic cell populations using the same approach, we observed moderate predicted differences in cell population frequencies as follows: >50% reduction in predicted skin/stratified epithelial cells (c14), >50% increase in osteoblasts (c11) and >40% increase in endothelial cells (c22) in mutant versus controls (Extended Data Fig. 5a). Comparison of gene expression differences between genotypes indicated extensive transcriptomic changes that are unique to non-hematopoietic cell populations (Extended Data Fig. 5a).

There were notable impacts on genes associated with hearing loss (e.g., collagens, *Notch1, Bmp4, Nfkb1, Bax, Egfr, Ptn Bmp4, Itgb1, Meis2*) and ossification (e.g., *Col1a1, Col2a1, Mef2c, Ctgf, Ecm1, Spp1*), both upregulated and down regulated (Extended Data Fig. 4b and Supplementary Table S8-S10). The CFTR centered, hearing associated and altered predicted Transcription Regulatory Networks (TRNs) are shown in Extended Data Fig. 4c. These effects were further evidenced from Gene Ontology, WikiPathways and Phenotype Ontology Enrichment analyses (Extended Data Fig. 5a-d and Supplementary Table S10, S11). It is critical for proper morphological alignment of middle ear ossicles for conduction and amplification of sound waves from the tympanic membrane to the inner ear. To check any genetic anomalies of CF middle ear ossicles, the expression of genes responsible for ossicle morphogenesis were evaluated (Extended Data Fig. 6). In the osteoblast cell population, *Tshz1, Dusp6, Fgfr1, Thra, Thrb* genes and, in the mesenchymal cell population *Tshz1, Thra and Thrb* genes are down regulated in the ΔF508 compared to WT.

Next, the transcriptional differences of middle ear epithelial cell types between WT and ΔF508 were examined. To this end, the epithelial cell cluster, based on their distinct expression of the epithelial cell adhesion gene, *Epcam* was exclusively selected. The significantly differentially expressed genes (DEGs) within the epithelial cell cluster were analyzed to identify the subsets with unique molecular characteristics. These correspond to the 5 initial ICGS2 identified epithelial populations, wherein CFTR expression was limited to the secretory 1 and secretory 2 subclusters (Fig. 4c and 4d). *Epcam+* subclusters were annotated based on the signature marker genes as follows: basal (DEGs: *Krt15*, *Lgals7, Dsc3);* secretory 1 (Sec1) (DEGs:*Reg3g, Muc5b*); secretory 2, (Sec 2) (DEGs: Reg3g, *Abi3bp, F3*); secretory 3 (Sec 3) (DEGs: *Cldn3, Kcnn4*); Ciliated (DEGs: *Sntn*, *Tmem212, Ccdc153, Dnah12*) (Fig. 4e).

A major imperfection in the cilia-generated bead flow was identified in ΔF508 mutant middle ear epithelial cells (Fig. 1l, 1m and 1n). Therefore, differentially expressed gene categories related to the ciliogenesis were investigated to identify, whether it was rooted to transcriptomic level (Fig. 4g). Expression of these gene categories were entirely restricted to the ciliated cell sub-cluster. They were further analyzed in depth using a manually curated list of ciliary development related transcripts^36^ to see to what extent gene expression patterns are altered in the ΔF508 sample. A broad deregulation of cilia associated Gene Ontology categories was displayed in the ΔF508 as compared to WT (Fig. 4g). Specifically, miscellaneous centrosome, centriole to cilium, axonemal assembly maintenance, ciliary trafficking, axonemal motility and miscellaneous cilia GO categories are broadly decreased in ΔF508. To understand how ΔF508 gene expression impacts alter cilia regulatory and signaling interactions, we constructed a network of protein-protein and putative regulatory interactions among cilia DEGs (Fig. 4h). In addition to downregulation of integrin associated singling components, we observe ΔF508 dependent downregulation of *Fam161a*, *Ift46* and *Ift172*, all factors required for cilia formation, function and/or maintenance^37–39^. These factors were connected through putative transcriptional regulation by *Stat6*, which was a non-regulated central node in this sub-network. Additional gene regulatory networks for the other epithelial cell subclusters, nominate alternative possible regulators of epithelial dysfunction, deregulated with ΔF508 (Extended Data Fig 7,8 and 9)

## Discussion

Over the past two decades, due to the discoveries of pharmacological treatments and changes in disease management practices, the life expectancy of children with CF has increased tremendously; the predicted life span for an individual with CF has grown to approximately 46 years (CFF patient registry, 2019). Therefore, it is essential to address CF-associated health issues such as hearing to improve quality of life. According to the National Health and Nutrition Examination Survey (NHANES), patients with cystic fibrosis have a significantly higher prevalence of hearing loss (31.8%) than that observed in persons without CF (14.9%). Early detection of hearing impairment is extremely important for speech and language development in childhood. Even a mild hearing impairment (16-40 dB) in children may be detrimental to speech development and communication^40,41^. Several studies have identified a higher prevalence of hearing loss in CF patients; this hearing loss is due to not only ototoxic effects that cause sensorineural hearing loss, but also due to conductive conditions that may be linked with otitis media and ET dysfunction^17,18,42–44^. Accumulating evidence supports that individuals with CF have a higher prevalence of developing otologic disorders, including otitis media^16,19,45,46^. Health of the middle ear is very likely related to CHL in patients with CF, yet the middle ear has not been extensively studied using physiologically relevant *in-vitro* models. Our study is the first to evaluate pathophysiological changes in the middle ear, and specifically to evaluate the role of CFTR in middle ear conductive hearing loss.

Many studies provide insight into the significance of ion and water balance/transport in the middle ear and the effect on hearing^47–49^. Sensorineural hearing, which is intimately related to the inner ear, is entirely regulated by ion transport. The process of transforming sound waves to neural signals, which includes depolarizing hair cells in response to sound waves, releasing neurotransmitters, and eliciting nerve signals to the brain, depends on CFTR^48^. Previously, *Choi. et al*., studied the ion channels critical for fluid homeostasis in the middle ear epithelium, including epithelial sodium channels (ENaC) and CFTR, using normal human middle ear epithelium (NHMEE) cultured in an ALI model^6^. They observed that CFTR expression is relatively less in NHMEEC than in the normal human nasal epithelium. Their prediction was that the human middle ear epithelium is adapted to fluid absorption rather than secretion.

The immunofluorescence imaging of our study confirmed the presence of CFTR protein expression on the apical side of the middle ear epithelium (Fig. 2c), highlighting its functional role in regulating apical membrane ion channels. However, the CFTR expression is limited to the Sec1 and Sec2 cell types. We developed an *ex-vivo* model using mouse auditory bulla connected to a perfusion system (Fig. 1b and 1c), which allowed measurement of CFTR function *in-situ*. The iodide efflux assay showed increased CFTR activity in WT compared to the ΔF508 mutant animals, confirming middle ear CFTR activity (Fig. 1e, 1f). We further validated our observation through the short circuit current assay in an ALI culture system (Fig. 1h, 1i).

Another important role of the middle ear epithelium appears to be eliminating cellular debris by muco-ciliary clearance and maintaining a fluid-free, air-filled cavity. Various simplified physical, mathematical and simulated middle ear models have been developed to reconstruct and to test middle ear functionality^50^. Most of these models assume that middle ear impedance is primarily due to “ossicular coupling” (transmission of sound waves through ossicles) ^51^. However, with inflammatory conditions of the middle ear, “acoustic coupling” (cochlear response due to pressure on oval and round window) plays a critical role in transmitting sound waves as well^51,52^. Consequently, inflammatory conditions of the middle ear must be addressed in studying conductive hearing loss, and multiple other mechanisms should be considered as contributory factors to CHL as well: decreased compliance of the TM, obstruction of ossicular chain movement, decreased sound pressure applied to the oval window-stapes footplate complex, or altered input of sound energy to oval and round windows^52^. Ultimately, a functional middle ear model is needed to comprehensively understand all mechanisms leading to CHL.

Wideband tympanometry has been used to assess the acoustic transfer function in various pathophysiological conditions to predict conductive hearing loss^53^. We obtained distinct Wide Band Absorbance (WBA) patterns under different epithelial conditions in the *ex-vivo* system; these patterns demonstrated the excellent sensitivity of our middle ear model. The WBA patterns indicated significantly reduced sound energy absorption at a high frequency range from 3000Hz-5000Hz and a discrete tympanogram pattern for ΔF508 cells compared to WT cells. Thus, our results indicate a deviation of acoustic transfer function related to CFTR dysfunction. As the ET, tympanic membrane, and the volume in the middle ear cavity are constants in our model, the observed change resulted entirely from the cellular condition of the ALI. However, additional studies are needed to understand how cellular phenotype (ΔF508) alters sound absorption and its correlation to CHL.

Recently, several studies evaluated gene expression in the middle ear through bulk RNA and scRNA analysis^54–57^. Our single cell transcriptomic analysis of middle ear cells generated 35 clusters, most of which included hematopoietic immune cells. The prominent cell clusters of immune cells are in both mutant and control groups. It has been reported that the innate immune response is acute and potent in the middle ear for rapid resolution of the otitis media^54^. Middle ear transcriptomic analysis detected the expression of 805 out of 809 innate immunity related genes (GO category 0045087). Taken together, these factors support the presence of an effective innate immune response in the middle ear.

Our study assessed the potential genetic components leading to CHL using the single cell transcriptomic profile of ΔF508 and WT middle ear. Major cell types between the two samples were very similar. However, these cell clusters showed notable differences in gene expression between ΔF508 mutant versus WT control group. These changes were more prominent in the gene expression profiles of hearing loss and ossification (Extended Data Fig. 5b). While most of the genes related to hearing and ossification were upregulated, genes involved in protein synthesis and extracellular matrix formation were inhibited in the ΔF508 mutant group (Extended Data Fig. 5b). The ΔF508 mutant epithelial cells showed higher gene expression levels related to cell proliferation, differentiation, migration, apoptotic inhibition, endothelial ossification, and collagen fibril structure formation.

It was well documented that the implication and distribution of cilia in the middle ear and ET^54^. These ciliated cell population is important for maintaining the mucus flow, which is vital for transferring pathogens from the tympanic cavity to nasopharynx through the auditory canal. We observed a significant decrease of cilia generated flow (Fig. 1n) and imperfect expression of genes related to ciliogenesis (Fig. 4g) in ΔF508 compared to WT, indicating a deficit mucociliary clearance. Similar ciliary anomalies and low ciliary beat frequency was observed in chronic otitis media^58^. Reduced epithelial secretions, condensed mucus and inflammatory mediators can diminished cilia function and decrease cilia generated flow.

In conclusion, this study confirmed the presence of CFTR expression in middle ear epithelium, its association with middle ear fluid secretion and altered cilia-generated flow in CF. Defective ciliogenesis, and dysfunctional ciliary action in the CF middle ear, may impact mucociliary clearance. We also described the impact of the middle ear epithelium on conductive hearing, and the extent of damage in the CF middle ear, with the middle ear model. We also described here the signature transcriptome level differences between WT and ΔF508 epithelium. Our findings indicate that increased expression of the genes related to inflammation in the middle ear epithelium and eustachian tube dysfunction due to defective cilia may play a role in CHL.

## Supporting information

Cilia generated bead flow of dF508 middle ear

Cilia generated bead flow of WT middle ear

Single cell RNA sequencing data

## ACKNOWLEDGEMENTS

None

## AUTHOR CONTRIBUTIONS

PL performed all the experiments unless specified, analyzed data, prepared the manuscript. APN supervised the study. PL, KSM, APN, KA designed the experiments. KSM fabricated the middle ear model. HLO performed cilia generated flow experiments. YH and YR assisted with animal husbandry. JL performed electrophysiological experiments. NS and GC performed single cell RNA sequencing data/bioinformatic analysis. LH carried out the tympanometry experiment. BRS, APN and KA assisted in editing the manuscript.

## ADDITIONAL INFORMATION

Supplementary table S1-S-11

Supplementary video 1

Supplementary video 2

## DATA AVAILABILITY

Sequence data that support the findings of this study have been deposited in the National Center for Biotechnology Information Gene Expression Omnibus ‘GenBank’ with accession code GSE226909 (reviewer token: ydmzscislfmhvkz).

## Methods

### Animals

All the animal procedures were conducted under protocols approved by Cincinnati Children’s Hospital Medical Center’s Institutional Animal Care and Use Committee, and in compliance with institutional and regulatory guideline. The WT and CFTR ^ΔF508/ΔF508^ mice (in-house colonies or provided by Craig Hodges, University of North Carolina, Chapel Hill, North Carolina, USA) were maintained in a barrier facility at CCHMC animal core. They were fed with normal chow, but the ΔF508/ΔF508 CFTR mice were treated with Colyte (osmotic laxative) water composed of polyethylene glycol 3350 (18 mM), NaCl (25 mM), KCl (10 mM), NaHCO_3_ (20 mM) and anhydrous Na_2_SO_4_ (40 mM). The protocol for isolation of mice auditory bulla, middle ear epithelial cells, culture and differentiation was followed as described previously ^22^.

### Mouse middle ear epithelial cell culture

Mouse middle ear epithelial cells isolation, culture and differentiation was carried out as described previously^22^. Following buffer solutions were made for this process.

1. mMEC basic media: penicillin (100 μg/mL) and streptomycin (100 μg/mL) in DMEM/F-12 HAMs media (Life Technology)
2. mMEC-FBS: mMEC basic with 10% BSA
3. mMEC+ media (proliferation media): 250 μL of the insulin (2 mg/ml stock), 50 μL of transferrin (5 mg/ml stock), 50 μL of the Cholera toxin (100 μg/ml stock), 250 μL of the EGF (5 μg/ml stock), 1.5 mg of BPE, 2.5 mL of FBS were added to 500 ml mMEC basic media. A 5 μL of Retinoic acid was added (5 mM stock) just before using the media.
4. mMEK-SF Media (differentiation media): 125 μL of the insulin (2 mg/ml stock), 50 μL of transferrin (5 mg/ml stock), 12.5 μL of the Cholera toxin (100 μg/ml stock), 50 μL of the EGF (5 μg/ml stock), 1.5 mg of the BPE, 500 μL of BSA to 49.1 mL of mMEC basic media.

Briefly, harvested middle ear cavities were immersed in pronase solution overnight at 4°C. In the next day, pronase was neutralized by adding 10% FBS. Then the samples were transferred to 2ml of fresh mMEC-FBS media, agitated the samples by inverting the tube for 25 times to dissociate cells. This process was repeated three times and combined the fractions containing dissociated cells. The solution was centrifuged at 500Xg for 5 min at 10°C. The cell pellet was resuspended in 5 ml of mMEC-FBS media. The cells were seeded at a density of 1 X10^4^ cells per transwell membrane in the 10 μM of Rho kinase inhibitor Y-27632 dihydrochloride. The cells were initially submerged in mMEC+ media with 300 μL of media in the top chamber and 700 μL in the bottom chamber. The media was changed every 48 hrs until the cells become fully confluent.

To differentiate the cell at air liquid interface (ALI) conditions, the media from top chamber was completely removed and 700 μL mMEC-SF media was added to the bottom chamber. The cells were fully differentiated in ∼ 14 days. The apical surface was washed with HBSS to remove the cellular secretions and mucus. These fully differentiated cells were used for Ussing chamber and middle ear on-a-chip experiments.

### Middle ear perfusion system

The peripheral tissues were carefully removed from the isolated middle ear cavities to see the clear bone at the outer ear opening and eustachian tube end. This task was performed very carefully, to not to damage auditory bulla. Then, tympanic membrane and ossicles were carefully removed under a stereomicroscope (Leica M165 FC, Leica Biosystems, Wetzlar, Germany). To perfuse buffer through the middle ear, peristaltic pump tubing (IDEX Health & Science, ISMATEC, Ref No 070535-031-ND, SC0051T) was connected to the outer ear opening (Fig 1b,1c) using 5 min epoxy (DEVCON).

After herding epoxy, PBS was perfused through the middle ear and checked for leaks. The buffer should enter from the outer ear opening and come out of the ET opening in a successfully sealed system (Fig. 1d) The auditory bulla was kept in PBS (on ice) until the perfusion system was made. The tube length at the buffer reservoir end was adjusted in which it takes ∼1 min to bring the buffer through tubing to the middle ear. The flow rate of the peristaltic pump was set for 20 μL/min.

### Iodide efflux (IE) assay

CFTR function of the middle ear epithelial cells was monitored using iodide efflux assay^59^. First, Sodium Iodide containing loading buffer (136 mM NaI prepared in buffer: 137 mM NaCl, 4.5 mM KH_2_PO4, 1 mM CaCl_2_, 1 mM MgCl_2_, 10 mM glucose, 5 mM HEPES, pH 7.2) was perfused to fill the middle ear cavity. After 20 min, the pump was run for 1 min to fil the middle ear with fresh loading buffer. Then the middle ear was incubated for another 40 min at room temperature (total 1hr). Then, Efflux buffer 136 mM NaI prepared in buffer: 137 mM NaNO_3_, 4.5 mM KH_2_PO4, 1 mM CaCl_2_, 1 mM MgCl_2_, 10 mM glucose, 5 mM HEPES, pH 7.2) was perfuse through the middle ear for 10 min to completely wash the loading buffer. This step is particularly important because traces of NaI will interfere with the final result. After washing step, aliquots were collected four times in one-minute intervals to establish a stable baseline. At the end of the fourth sample collection, immediately switched to the flux buffer with CFTR agonists (Forskolin (10 μM) and IBMX (100 μM)) and aliquots were collected ten times at 1 min intervals. The iodide concentration of all 15 aliquots collected was determined using an iodide-selective electrode (Thermo Scientific, Waltham, MA) and converted to iodide efflux rate (nmol/min) as described previously^60^.

### Short-circuit current measurement

Polarized mice meddle ear epithelial cells cultured on Transwell support were mounted to an Ussing chamber system (Physiologic Instruments). The system was maintained at 37°C. Middle ear cells bathed in Ringer’s solution (mM) (basolateral side: 140 NaCl, 5 KCl, 0.36 K_2_HPO4, 0.44 KH_2_PO4, 1.3 CaCl_2_, 0.5 MgCl_2_, 4.2 NaHCO_3_, 10 HEPES, 10 glucose, pH 7.2, [Cl^-^] = 149), and low Cl^-^ Ringer’s solution (mM) (apical side: 133.3 Na-gluconate, 2.5 NaCl, 0.36 K_2_HPO_4_, 0.44 KH_2_PO_4_, 5.7 CaCl_2_, 0.5 MgCl_2_, 4.2 NaHCO_3_, 10 HEPES, 10 mannitol, pH 7.2, [Cl–] = 14.8) and cells were supplied with 95% O_2_ and 5% CO_2_. First, cells were treated with amiloride (50 μM), after current stabilization, CFTR was activated by adding forskolin (10 μM) and IBMX (100 μM). To verify the CFTR dependency of currents, CFTR_inh_172 (20 μM) was added to the apical side of the cells.

### Cilia generated bead flow measurement

Cilia generated bead flow for middle ear epithelial cells were performed as describe previously^61^. Briefly, after isolation of auditory bulla, it was segmented into smaller pieces under stereomicroscope (Leica M165 FC, Leica Biosystems, Wetzlar, Germany). These pieces were kept in PBS and and mounted in 1-mm-spaced slice anchors (Warner Instruments, Hamden, CT, USA). The middle ear fragmets were placed under BX61WI fixed-stage motorized upright microscope (Olympus, Tokyo, Japan). FITC-conjugated beads (0.5 mm) were added to the samples. Bead flow was measured at 32 frames/s. Quantitation of bead velocity was performed as described earlier^62^.

### Fabrication of a novel *in-vitro* middle ear model (middle ear on-a-chip) for hearing test

The hearing test device contains eardrum-like structure and Eustachian tube-like structure created by pouring polydimethylsiloxane (PDMS; Ells Worth Adhesive; # 4019862) on a mold that was designed in house and created by a 3D printer. The PDMS was prepared by mixing thoroughly with curing kit at a ratio of 10:1 (wt%) and degassed in a desiccator (Fisher Scientific; #50-212-719) under vacuum system for 40 min. In the meantime, the mold placed in a 24-well plate (costar, #3527) and was coated with trichloro silane (Sigma-Aldrich; #448931) in another desiccator under vacuum system for 30 min. The silane coating helps to separate the PDMS device from the mold. Degassed PDMS was casted onto the mold using a 1 mL pipette with a tip.

The end of the tip was cut using a scissors to make wide-end due to the uncured PDMS is very viscous. The PDMS layer was separated from the mold following cure at 60 °C for at least 4 hours. Eardrum-like membrane was created with a highly elasticity silicon rubber, Room Temperature Vulcanized 615 (RTV 615; Momentive; #9480) ^63^, as a flexible thin layer of membrane. Uncured RTV 615 was mixed with curing kit at a ratio of 5:1 (wt%) and degassed in a desiccator for 30 min and a 3’’ polished-flat silicon wafer was coated with trichloro silane in a desiccator for 30 min as before. The silane-coated silicon wafer was placed on a vacuum chuck in a spin coater (Specialty Coating Systems; #6800) and spun by a following programming for 5 µm thickness of eardrum-like membrane after pure degassed RTV 615 on the wafer: (1) ramp up to 500 rpm for 10 sec and holding the speed for 10 sec; (2) ramp up to 3000 rpm for 10 sec and holding the speed for 5 min; (3) speed down to 0 rpm for 10 sec. It placed on a hot plate (Fisher Scientific; #HP88857204) at 120 °C for 1 h and was cool down to room temperature. We assembled the thin layer of eardrum-like membrane with the *in vitro* hearing test device followed by treatment of the two surfaces, which are contacting area, with oxygen plasma for 30 sec using a Tergeo Plasma Cleaner (PIE Scientific).

After exposed the two surfaces, put the PDMS device down on the plasma-treated RTV 615 silicon rubber on the wafer and placed them on the hot plate at 120 °C for 20 min and turned off the power to cool down to room temperature. After cooling down, the PDMS device with the thin layer of membrane was peeled off from the silicon wafer and cut off unexpected thin layer of the membrane using a sharp knife. Eustacian tube-like structure was created using a needle (BD; #305106; 30G) by poking from an edge of surface to inner-extruded tunnel (Supplementary Figure) to release pressure in the space between the eardrum like membrane and cell membrane.

### Tympanometry

To test the middle ear on-a-chip, middle ear cells were isolated and cultured on Transwell insert (Corning; #3470) and differentiated under air-liquid interface (ALI) conditions as describe previously^22^. These cells were maintained in an incubator at 37°C, 5% CO_2_ until they are used for the test. Right before taking the tympanometric measurement, Transwell inserts were taken out of the incubator and the top hanging support was removed using a diagonal cutting plier. Then, it was inserted into the 3D printed middle ear modal. The function of the middle ear-chip was measured using wideband tympanometry using a click stimulus (Interacoustics Titan wideband software and hardware) with absorbance measured as a function of frequency from 250-8000 Hz. The ratio of the absorbance, calibrated in a 2-cc coupler relative to the absorbance in situ was measured. After assembling the middle ear chip, the tip of a tympanometer probe was connected using a silicone sleeve that matched the diameter of the simulated outer ear canal, and measurements are taken 3 successive times in each condition. The repeated measures were averaged and the mean plus one standard deviation are shown in Fig 3c and 3d.

### RNAScope

Formalin fixed, paraffin embedded (FFPE sections), decalcified, middle ears were sectioned to see both middle ear cavity and eustachian tube (Fig 2a, Extended Data Fig 1). The slides containing these middle ear sections were used for RNAscope assays. RNAScope was performed using the Multiplex Fluorescent Reagent Kit V2 Assay (Advanced Cell Diagnostics, Inc.) according to the manufacturer’s user manual. All the RNAscope probes were purchased from ACDBio. Mouse positive control probe (Cat No 320881) and negative control probe (Cat No320871) were used. The CFTR transcripts in the middle ear FFPE sections were analyzed by the mCFTR probe (Cat no 483011). To compare the CFTR expression levels in the middle ear, highly abundant Muc5b (Cat No 471991) transcripts were analyzed.

### Immunofluorescence

Mouse middle ear sections containing slides blocked at room temperature for 1 hour in Tris-buffered saline with 1% BSA, 5% normal goat serum, and incubated in primary antibody, mouse anti-CFTR 3194 anti-body (obtained from Dr. Marino, Memphis, TN, which has been extensively validated) overnight at 4°C, followed by wash and secondary antibody (Alexa Fluor 568, Invitrogen; 1:500) incubation for 1 hour. Slides were washed and mounted with prolong gold antifade reagent. Images were acquired on the Olympus Inverted Confocal Microscope (Olympus FV1200) with 60× objective and laser settings adjusted using positive and negative controls. All subsequent images were acquired with a fixed optical/exposure configuration.

### Mouse middle ear cells isolation for single cell RNA seq analysis

Age matched adult mice were used for middle ear isolation. Extracted mouse middle ear tissues were placed in ice-cold hypothermosol (Sigma, Cat# H4416) on ice. The tissues were transferred to a petri dish on ice and were minced for 3-4 min on ice into 1-mm’ pieces using a razor blade. The minced tissues were placed in 1 ml Bacillus Licheniformis Enzyme cocktail (100 μL b. lich 100 mg/mL (10 mg/mL final conc. - Sigma, P5380), 1 μL 0.5 M EDTA (0.5 mM final conc. - Sigma, A8806), 899 μL DPBS (no Ca, Mg) ThermoFisher (cat. #14190)) and incubate on ice. The sample was shaken every minute and triturated every 2 min using a p1000 pipet tip. After 20 min, the sample was transferred into a dounce homogenizer (10 strokes of Pestle, every 2 min for total of 8 min). The supernatant with released cells was transferred through a 30 μM filter to a 15 ml tube. The filter was rinsed with 5 ml of ice-cold PBS/0.04% BSA buffer solution. The cells were centrifuged for 300 g for 5 min at 4°C. The supernatant was removed, and cells were resuspended 100 μL ice-cold PBS/0.04% BSA buffer solution. 1 ml of Red Blood Cell Lysing Buffer (Sigma, cat # R7757) was added to the cells and incubated for 2 min on ice. Then 12 ml of ice-cold PBS/0.04% BSA buffer solution was added and centrifuged 300 g for 5 min. Cells were examined by hemocytometer with trypan blue. Cell concentration was adjusted to 1000 cells/μL for 10X Chromium.

### Single-cell RNA-Seq analysis

scRNA-Seq 10x Genomics FASTQ files were aligned to the mouse genome (mm10) using the Cell Ranger workflow (version 3.1.0) and the mm10-2.1.0 reference transcriptome. Each sample was sequenced to a depth of >340 million reads, with over 7,200 called cell barcodes from each library, with >46,000 reads per cell on average (default filtering options) (see summary metrics in Table S1). Using the default filtered cellular barcodes, the associated sparse-filtered HDF5 files were jointly processed (WT and ΔF508) in AltAnalyze prior to unsupervised clustering. Evidenced sex-specific transcripts (n=315, Table S2) were excluded from the scaled count matrix prior further analyses to limit any sex-based cluster or differential expression changes, as detailed below. ICGS2 (Iterative Clustering and Guide-gene selection version 2) to obtain normalized gene counts (counts per ten thousands (CPTT) UMIs) and initial predicted cell populations^65^. ICGS2 was run using the default options (cosine clustering) in addition to a PageRank down-sampling threshold of 5,000 cells. ICGS2 identifies only transcriptionally stable cell populations, that have multiple unique marker genes and restricted to only those cells that can be confidently re-assigned back to each cluster through supervised machine learning classification. Cell types were initially predicted based on ICGS2’s single-cell marker gene database of over 2,000 prior study mouse cell-type signatures and refined based on the literature or ToppCell enrichment. Comprehensive differential expression and gene regulatory prediction analyses were performed using these alignment results in the software cellHarmony with the supplied ICGS2 groups for both WT and ΔF508 (options --referenceType None) and eBayes t-test p<0.05^66^. Pathway and Gene Ontology over-representation analyses and associated heatmaps were produced using the GO-Elite module of AltAnalyze. Included UMAP plots were produced by ICGS2 or by AltAnalyze by querying the expression of individual genes (--genes “<GENE symbols>” --separateGenePlots True). The single-cell genomics data has been deposited in the Gene Expression Omnibus database (GSE226909; reviewer token: ydmzscislfmhvkz). Protein-interaction networks were produced in AltAnalyze using the NetPerspective workflow, for cellHarmony differentially expressed genes alone or with genes interest (e.g., *Cftr*).

To identify sex-associated transcripts, we combined evidence from two prior studies. Epithelial associated sex-biased genes were obtained from the prior single-cell kidney analysis^67^, while hematopoietic sex-biased genes were identified by identifying differentially expressed genes in bone marrow progenitors from male versus female mouse littermates profiled using the 10x Genomics platform across 20 cell populations (identified in at least 2 cell-populations, fold > 1.2 and eBayes t-test p<0.05, FDR)^64^.

### Statistical analysis

Data was generated from at least three independent replicates. The level of marginal significance, p-value, was calculated using two-tailed Student’s t test for pairwise comparison and one-way analysis of variance with Bonferroni adjustment for multiple variations. A p value <0.05 was considered significant.

---

**Extended Data Fig. 1.**
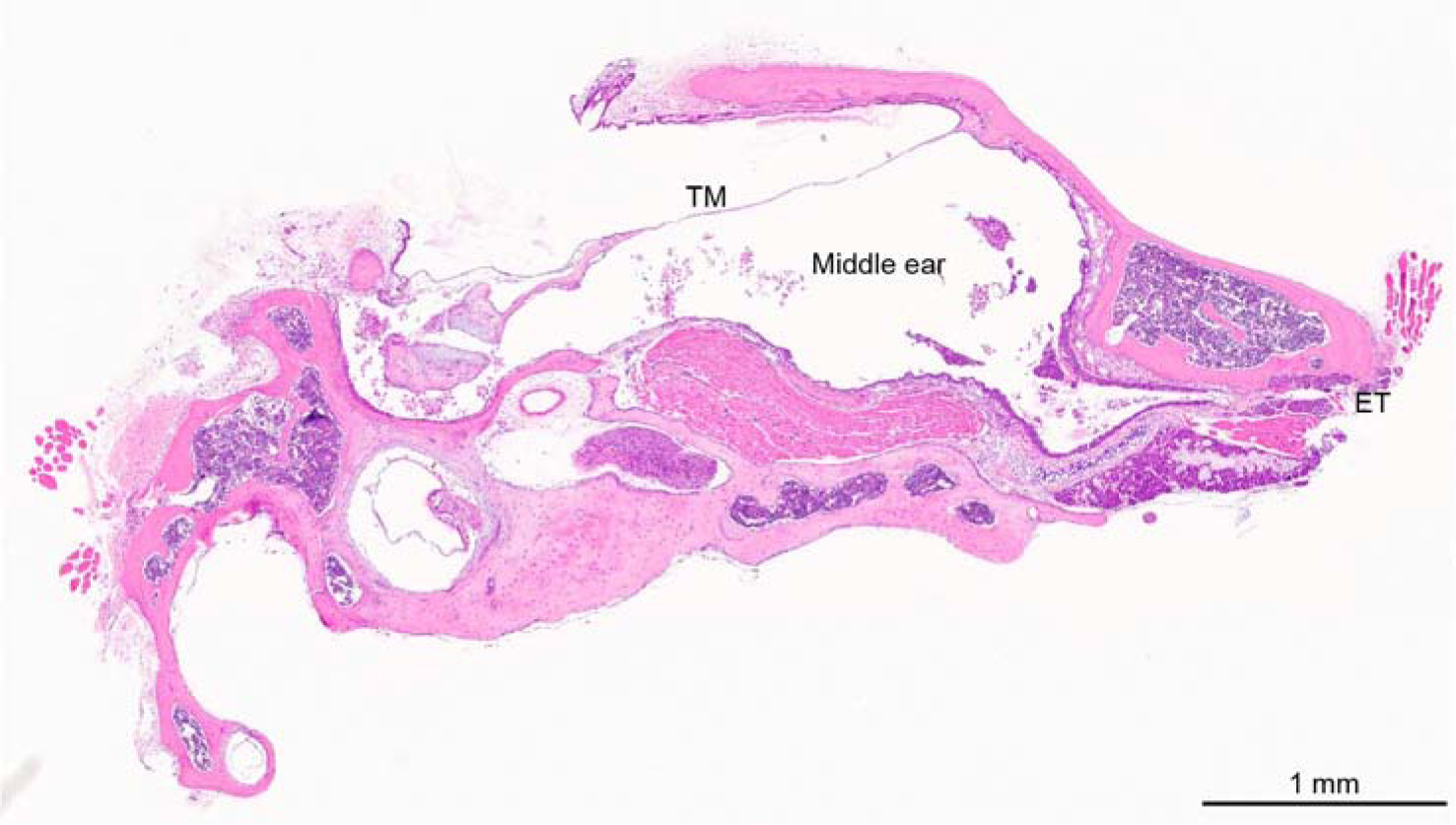
A cross-section of an H&E stained ΔF508 mouse middle ear. Image illustrates the positions of Eustachian tube (ET), middle ear cavity and tympanic membrane (TM).

**Extended Data Fig. 2.**
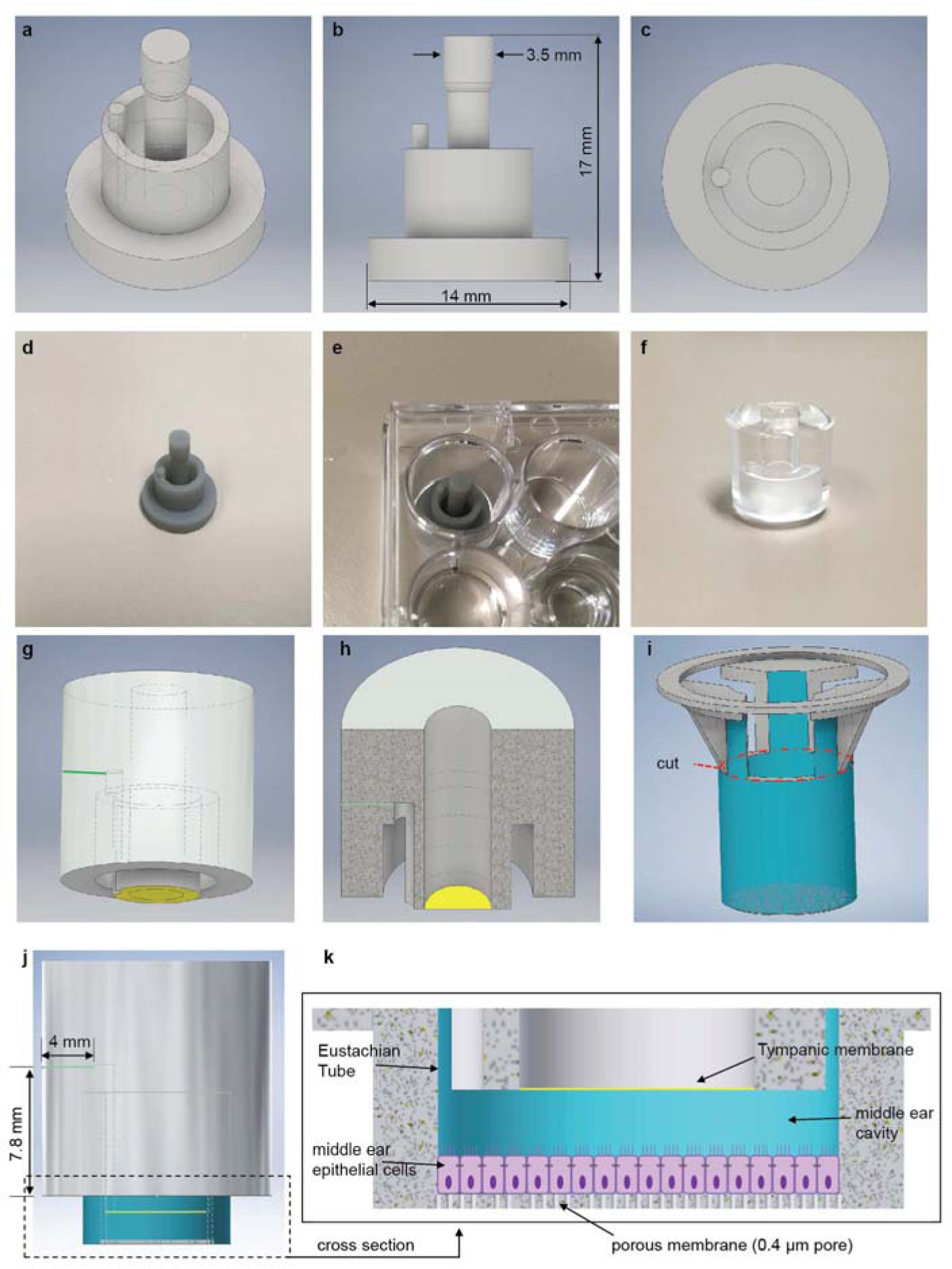
Fabrication of *in-vitro* middle ear on-a-chip hearing test model using a 3D printer. **a-d,** The mold used to make the middle ear modal was designed in house; **a**, isometry view **b**, front view and **c**, top view **d**, the 3D printed mold. **e**, The mold was placed in a 24-well plate and coated with trichloro silane. The hearing test device was fabricated by casting PDMS and curing at 60°C for overnight. **f**, The fabricated model after PDMS casting. **g**, A schematic of the fabricated hearing test modal. The attached tympanic membrane is shown in yellow. **h**, A schematic of a cross section of the fabricated hearing test modal. **i**, A schematic of a Transwell insert. Middle ear epithelial cells are grown on the Transwell membrane. It was cut along the red line to insert into the middle ear modal. **j**, a schematic of a trimmed Transwell insert incorporated middle ear modal. **k**, an enlarged cross section of “j” showing the positions of middle ear cavity, epithelial cells, tympanic membrane, eustachian tube.

**Extended Data Fig 3.**
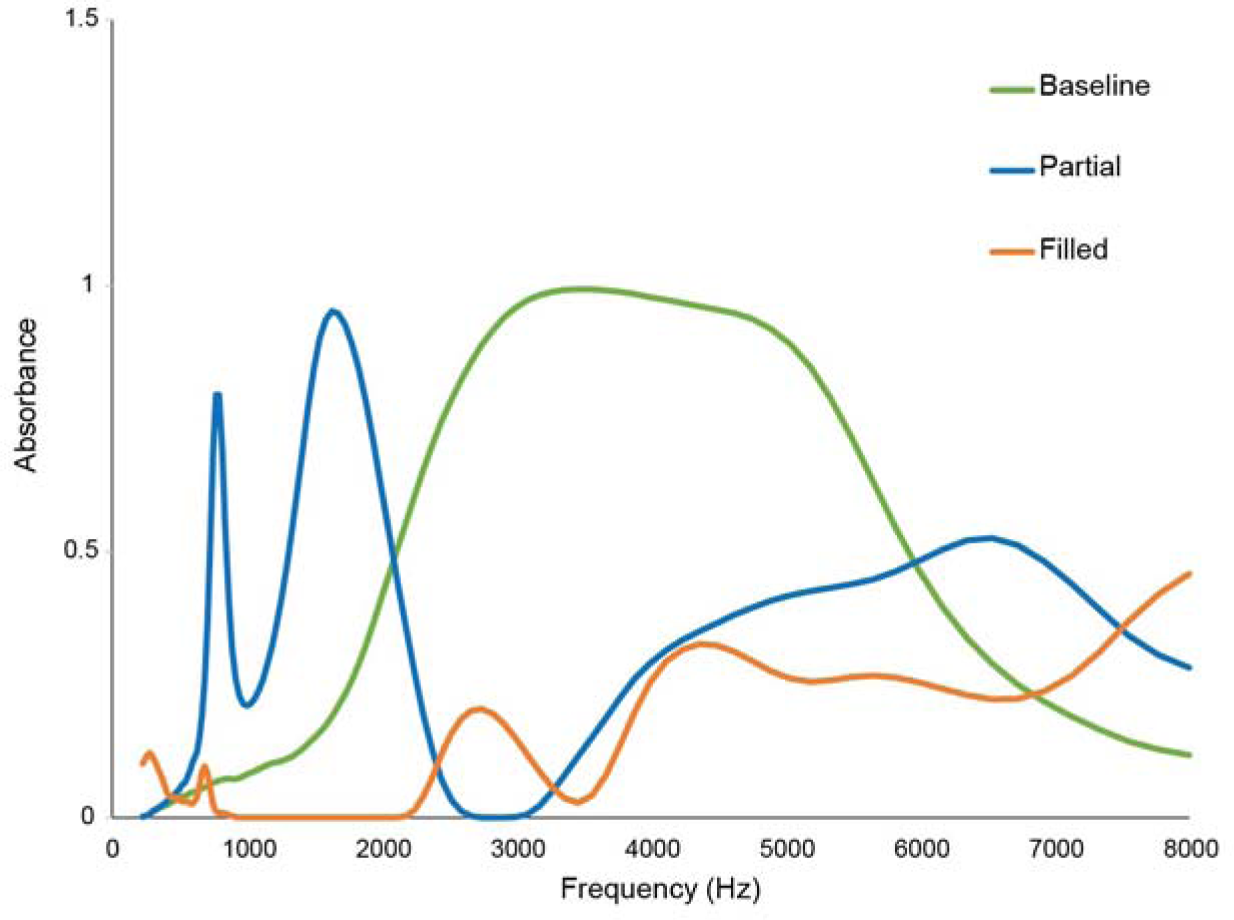
Wide Band Tympanometry measurement were taken with the fluid filled middle ear-chip. The absorbance results of the partial and fully filled states of the middle ear on-a-chip akin to the otitis media condition in real middle ear.

**Extended Data Fig. 4.**
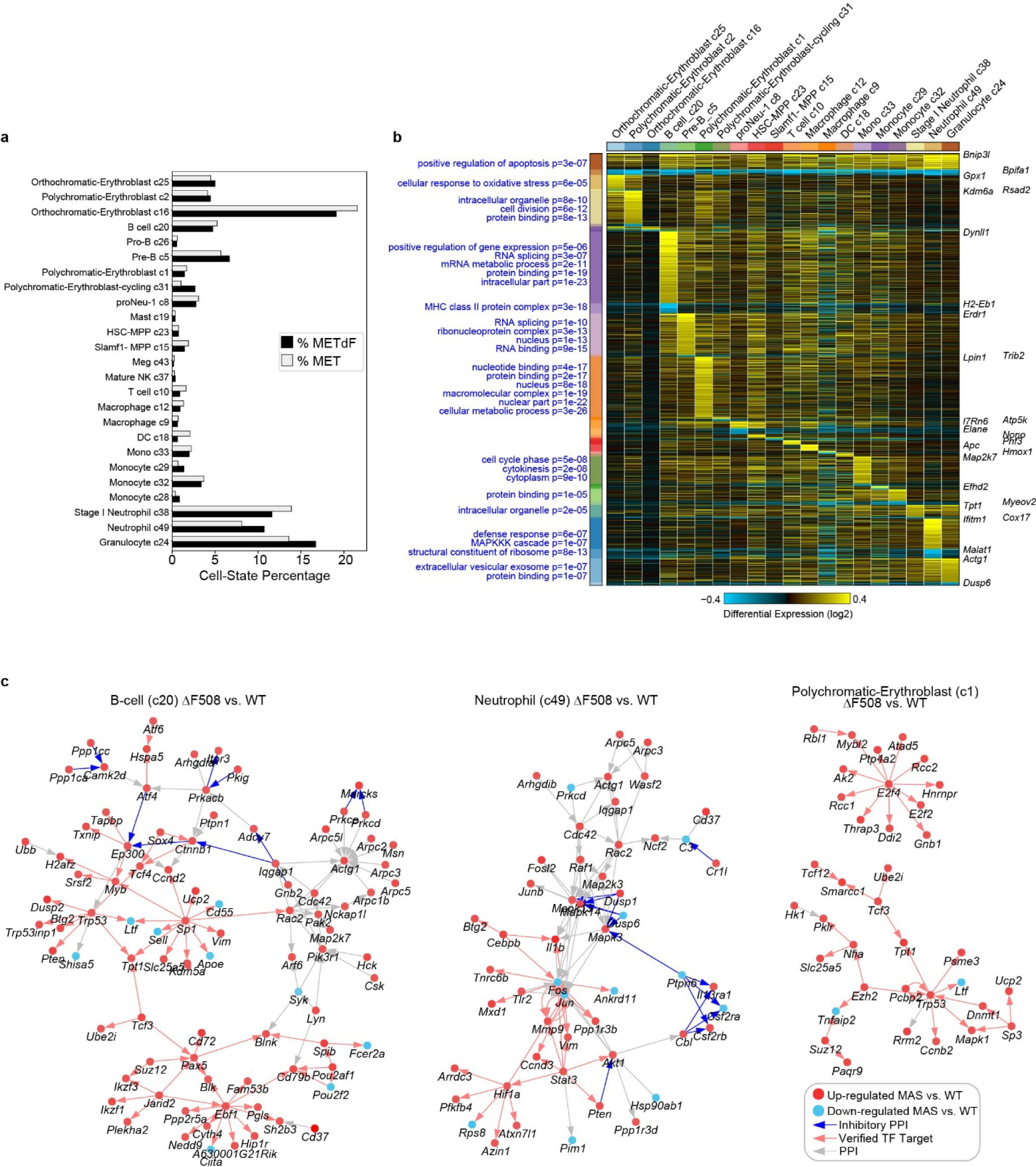
Population-specific impacts to distinct haemopoietic cell lineages with ΔF508 *Ctfr*. **a,** The percent of each hematopoietic cell population from the combined scRNA-Seq analysis shown in mutant relative to wild-type captures. **b,** Heatmap of ΔF508 versus control fold-changes (log2) for all differentially expressed genes comparing the same hematopoietic cell populations detected and organized by the software cellHarmony. The heatmap is broken down according to broad and cell-type specific patterns of F508 induced gene changes, with enriched Gene Ontology terms displayed to the left of the heatmap with the associated GO-Elite enrichment p-values. To the right of the heatmap are example gene callouts from each cellHarmony organized fold-change cluster. **c,** cellHarmony predicted gene-regulatory networks for the indicated cell-type comparisons (ΔF508 versus WT). See the in-figure legend for node and edge definition.

**Extended Data Fig 5.**
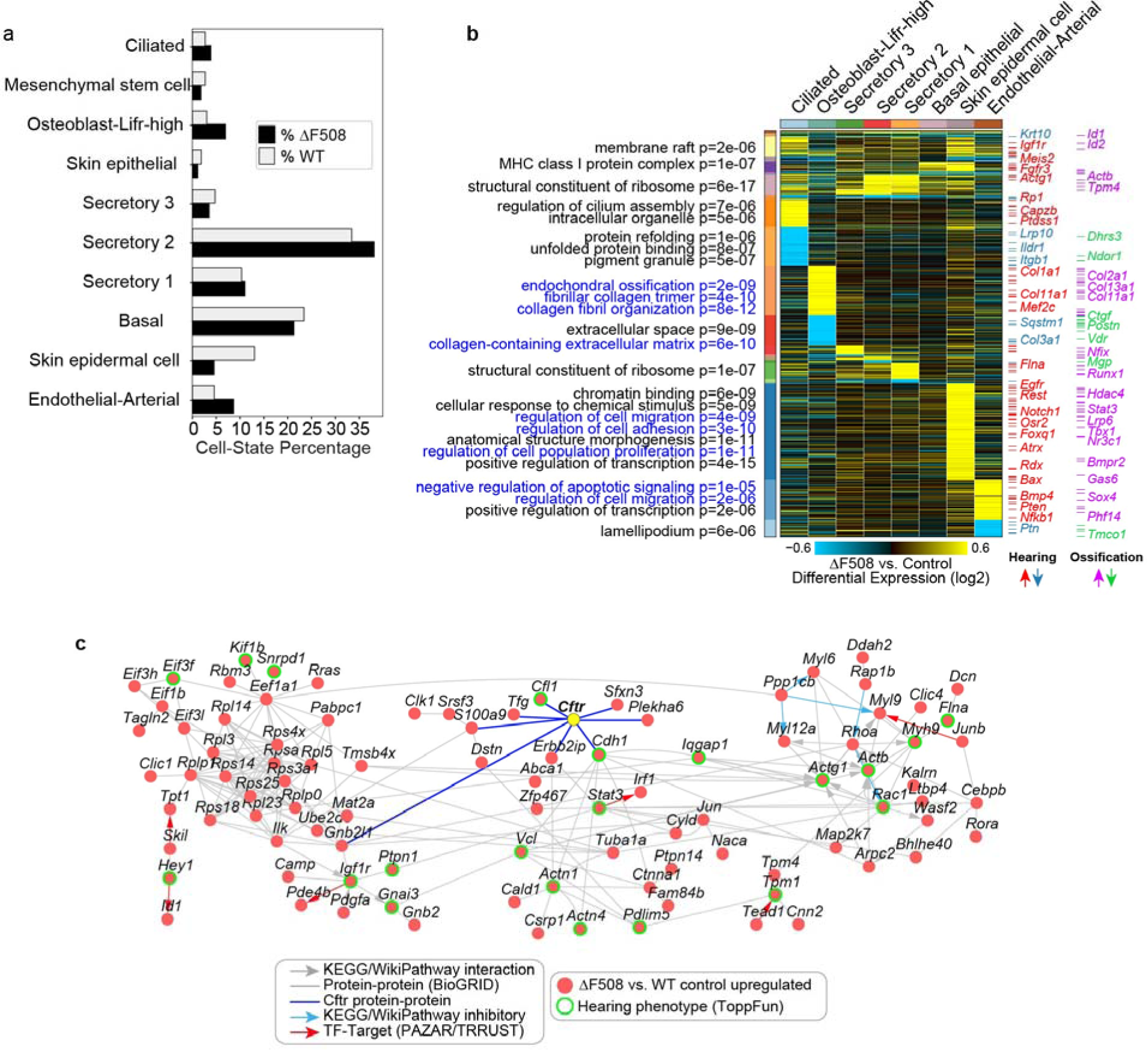
The differentially expressed genes (DEGs) associated with hearing loss and ossification shows a notable impact on ΔF508 middle ear. **a,** The percentages of each non-hematopoietic cell population is shown in mutant relative to wild-type captures. **b,** Heatmap of ΔF508 versus control fold-changes (log2) for all differentially expressed genes comparing the same non-hematopoietic cell populations detected and organized by the software cellHarmony. The heatmap is broken down according to broad and cell-type specific patterns of ΔF508 induced gene changes, with enriched Gene Ontology terms displayed to the left of the heatmap with the associated GO-Elite enrichment p-values. To the right of the heatmap are annotated genes (tick marks) associated with ossification and hearing identified from the software ToppGene, with example gene callouts from each category (colored by the indicate direction of regulation). **c,** The putative transcription regulatory networks that are directly interacting with CFTR in relation to hearing.

**Extended Data Fig. 6.**
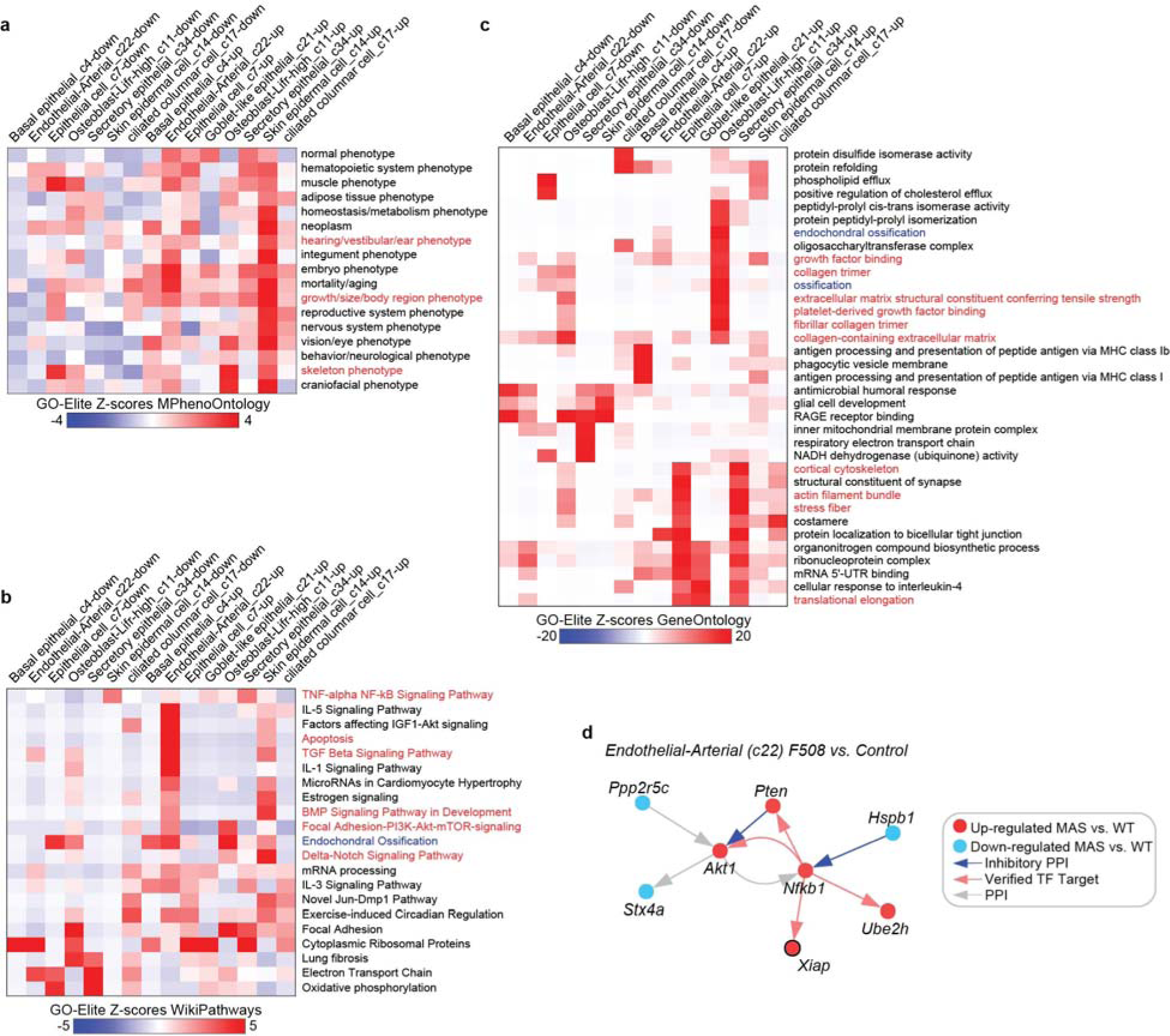
Perturbed core regulatory networks of. Δ**F508 *Ctfr* in the middle ear. a-c,** GO-Elite enrichment results for **a,** mouse phenotype ontology, **b,** WikiPathways and **c,** Gene Ontology gene-sets. Hearing loss-disease relevant terms are highlighted in red (hearing) or blue (ossification). **d,** Endothelial cellHarmony predicted core gene regulatory network.

**Extended Data Fig. 7.**
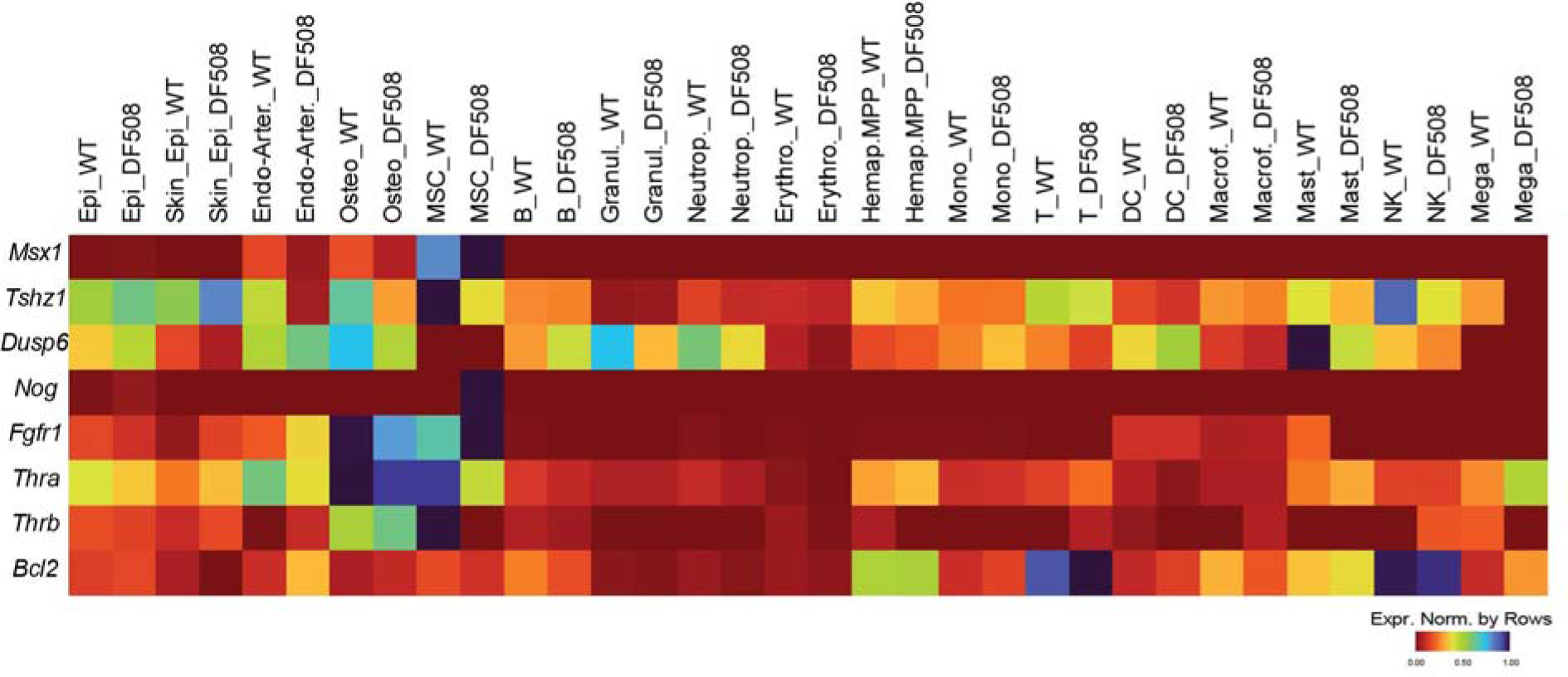
Differential expression of genes important for ossicle morphogenesis. Genes in the osteoblast and mesenchymal cell population are down regulated in the ΔF508 compared to WT.

**Extended Data Fig. 8.**
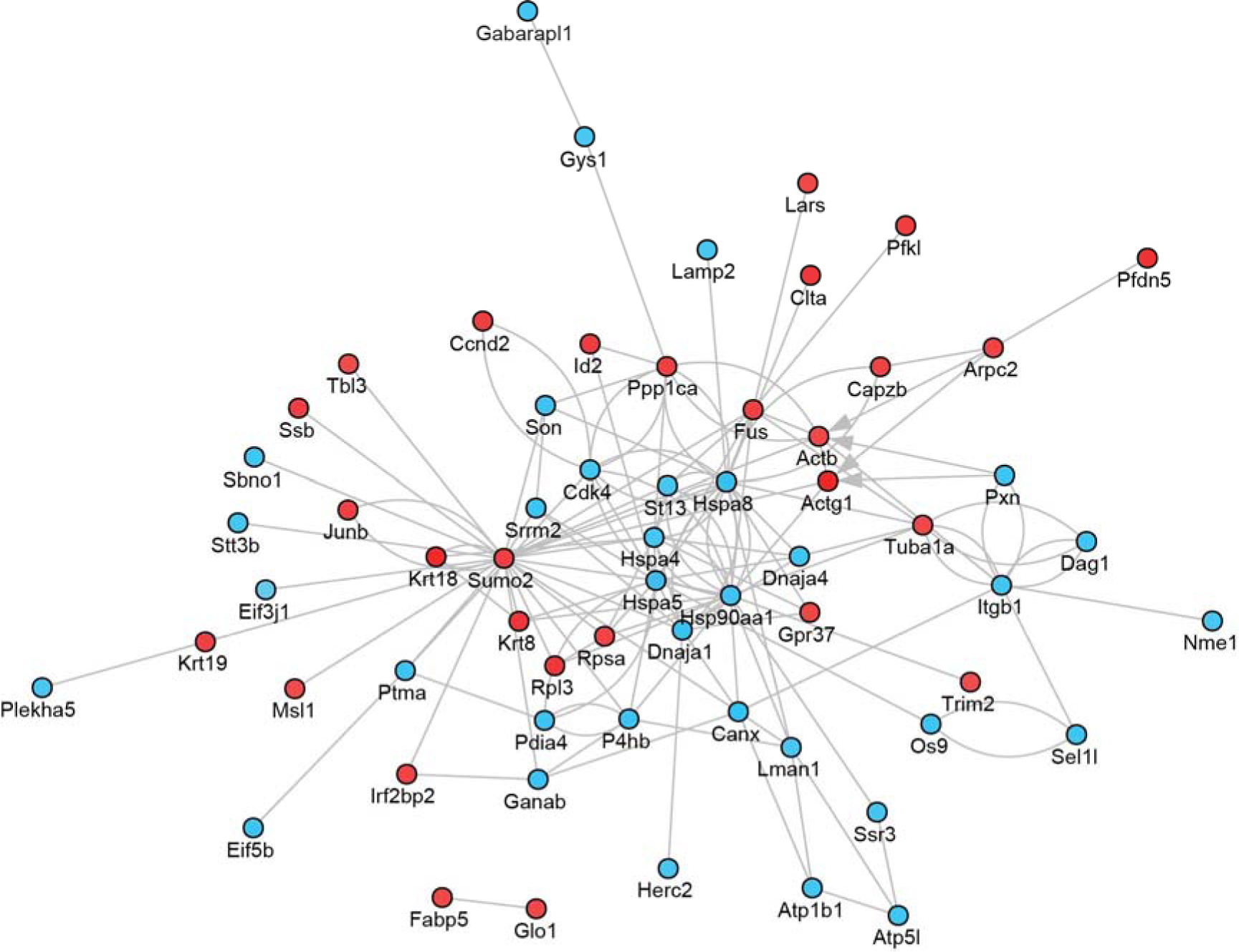
CellHarmony predicted core gene regulatory networks in ciliated cell population. Red dot indicates the down regulated genes and blue indicates upregulated genes.

**Extended Data Fig. 9.**
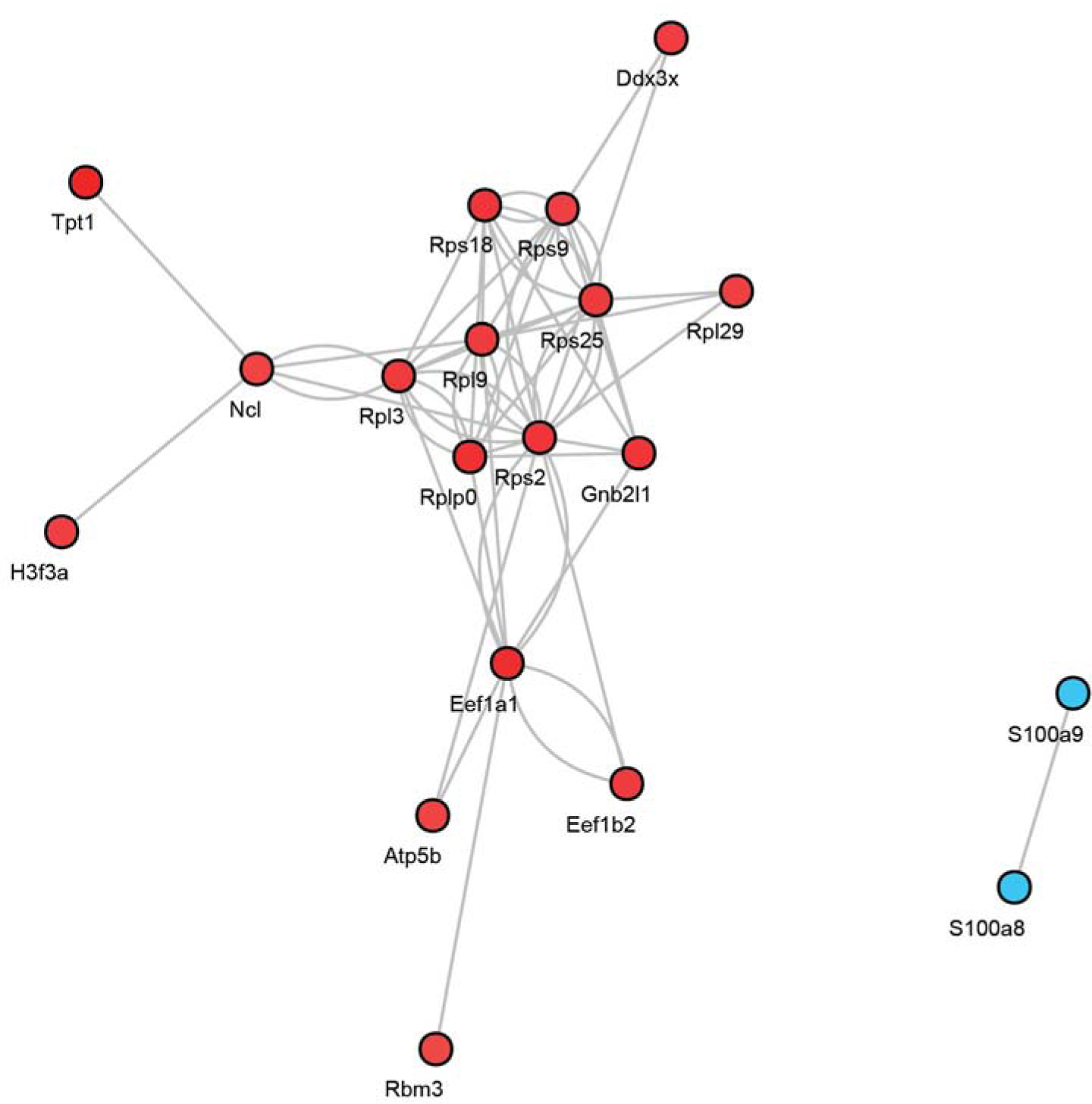
Transcription regulatory networks in goblet like secretory cell population. Red dot indicates the down regulated genes and blue indicates upregulated genes.

**Extended Data Fig. 10.**
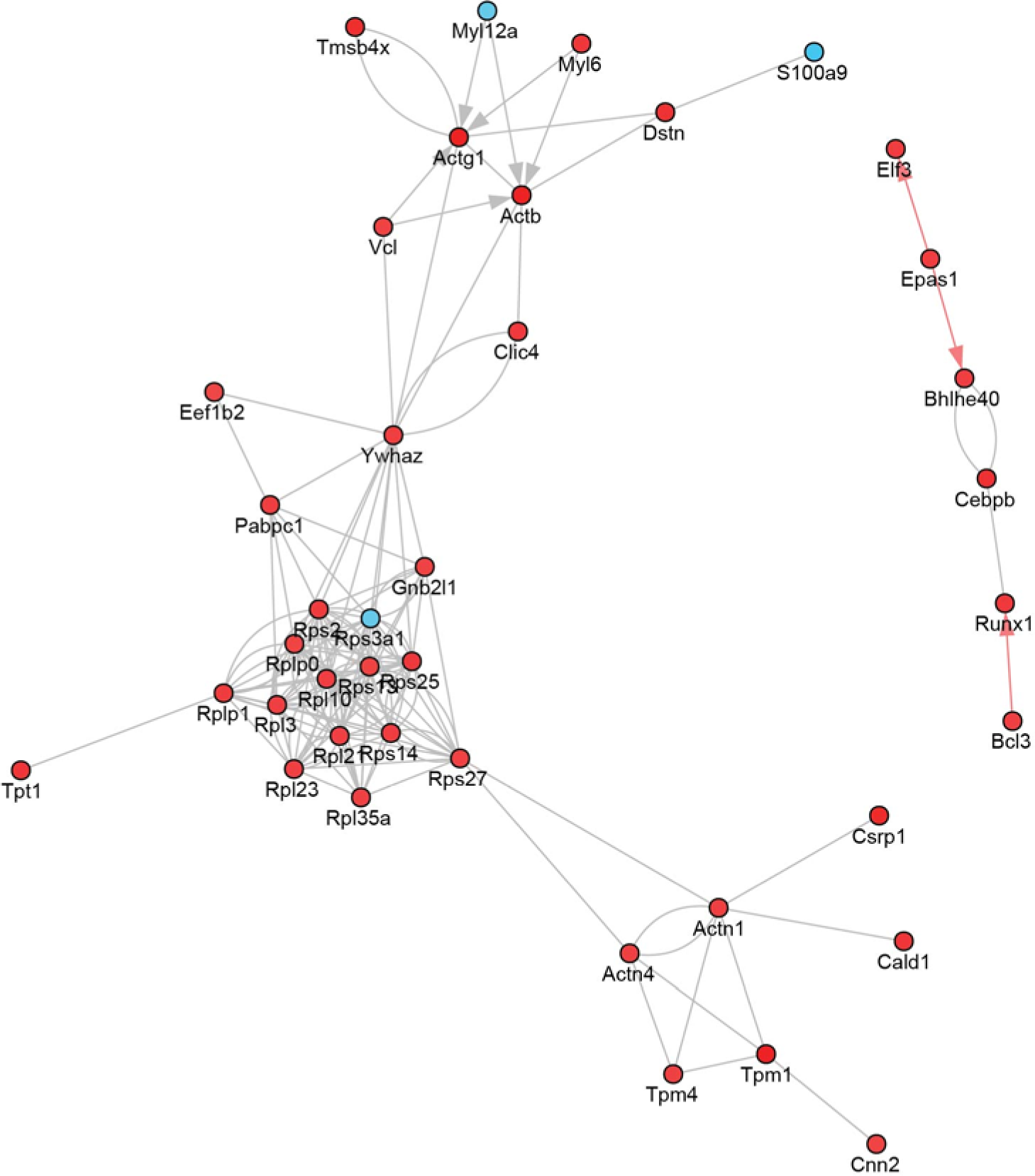
CellHarmony predicted core gene regulatory networks in secretory epithelial cell population. Most of the genes in ΔF508 sample are down regulated indicating the impact on secretory epithelial cell population. Red dot indicates the down regulated genes and blue indicates upregulated genes.

## References

1 Pfaff, C., Schultz, J. A. & Schellhorn, R. The vertebrate middle and inner ear: A short overview. J Morphol 280, 1098–1105 (2019).

2 Cunsolo, E. et al. Functional anatomy of the Eustachian tube. Int J Immunopathol Pharmacol 23, 4–7 (2010).

3 Juhn, S. K. et al. Determining otitis media severity from middle ear fluid analysis. Ann Otol Rhinol Laryngol Suppl 163, 43–45 (1994).

4 Zahnert, T. The differential diagnosis of hearing loss. Dtsch Arztebl Int 108, 433–443; quiz 444 (2011).

5 Morris, L. M. et al. Mouse middle ear ion homeostasis channels and intercellular junctions. PLoS One 7, e39004 (2012).

6 Choi, J. Y. et al. ENaC- and CFTR-dependent ion and fluid transport in human middle ear epithelial cells. Hear Res 211, 26–32 (2006).

7 Herman, P. et al. Ion transports in the middle ear epithelium. Kidney Int Suppl 65, S94–97 (1998).

8 Inagaki, M. et al. Ultrastructure of mucous blanket in otitis media with effusion. Ann Otol Rhinol Laryngol 97, 313–317 (1988).

9 MacArthur, C. J., Hausman, F., Kempton, J. B. & Trune, D. R. Murine middle ear inflammation and ion homeostasis gene expression. Otol Neurotol 32, 508–515 (2011).

10 Trune, D. R. Ion homeostasis in the ear: mechanisms, maladies, and management. Curr Opin Otolaryngol Head Neck Surg 18, 413–419 (2010).

11 Kerem, B. et al. Identification of the cystic fibrosis gene: genetic analysis. Science 245, 1073–1080 (1989).

12 Riordan, J. R. et al. Identification of the cystic fibrosis gene: cloning and characterization of complementary DNA. Science 245, 1066–1073 (1989).

13 Cheng, S. H. et al. Defective intracellular transport and processing of CFTR is the molecular basis of most cystic fibrosis. Cell 63, 827–834 (1990).

14 Marson, F. A. L., Bertuzzo, C. S. & Ribeiro, J. D. Classification of CFTR mutation classes. Lancet Respir Med 4, e37–e38 (2016).

15 Jorissen, M., De Boeck, K. & Feenstra, L. Middle ear disease in cystic fibrosis. Int J Pediatr Otorhinolaryngol 43, 123–128 (1998).

16 Bak-Pedersen, K. & Larsen, P. K. Inflammatory middle ear diseases in patients with cystic fibrosis. Acta Otolaryngol Suppl 360, 138–140 (1979).

17 Kreicher, K. L. et al. Audiometric assessment of pediatric patients with cystic fibrosis. J Cyst Fibros 17, 383–390 (2018).

18 Blankenship, C. M. et al. Functional Impacts of Aminoglycoside Treatment on Speech Perception and Extended High-Frequency Hearing Loss in a Pediatric Cystic Fibrosis Cohort. Am J Audiol 30, 834–853 (2021).

19 McCoy, J. L., Kaffenberger, T. M., Yang, T. S. & Dohar, J. E. Otitis media prone children with cystic fibrosis: A new normal. Am J Otolaryngol 42, 103137 (2021).

20 Thornton, J. L. et al. Conductive hearing loss induced by experimental middle-ear effusion in a chinchilla model reveals impaired tympanic membrane-coupled ossicular chain movement. Journal of the Association for Research in Otolaryngology 14, 451–464 (2013).

21 Shik Mun, K., et al. Patient-derived pancreas-on-a-chip to model cystic fibrosis-related disorders. Nat Commun 10, 3124 (2019).

22 Mulay, A., Akram, K., Bingle, L. & Bingle, C. D. Isolation and Culture of Primary Mouse Middle Ear Epithelial Cells. Methods Mol Biol 1940, 157–168 (2019).

23 Chen, Y. et al. Human primary middle ear epithelial cell culture: A novel in vitro model to study otitis media. Laryngoscope Investig Otolaryngol 4, 663–672 (2019).

24 Mulay, A. et al. An in vitro model of murine middle ear epithelium. Dis Model Mech 9, 1405–1417 (2016).

25 Luo, W. et al. Cilia distribution and polarity in the epithelial lining of the mouse middle ear cavity. Sci Rep 7, 45870 (2017).

26 Sade, J. Middle ear mucosa. Arch Otolaryngol 84, 137–143 (1966).

27 Thompson, H. & Tucker, A. S. Dual origin of the epithelium of the mammalian middle ear. Science 339, 1453–1456 (2013).

28 Tucker, A. S. et al. Mapping the distribution of stem/progenitor cells across the mouse middle ear during homeostasis and inflammation. Development 145 (2018).

29 Collawn, J. F. & Matalon, S. CFTR and lung homeostasis. Am J Physiol Lung Cell Mol Physiol 307, L917–923 (2014).

30 Marino, C. R., Matovcik, L. M., Gorelick, F. S. & Cohn, J. A. Localization of the cystic fibrosis transmembrane conductance regulator in pancreas. J Clin Invest 88, 712–716 (1991).

31 Bergeron, C. & Cantin, A. M. Cystic Fibrosis: Pathophysiology of Lung Disease. Semin Respir Crit Care Med 40, 715–726 (2019).

32 Durie, P. R. The pathophysiology of the pancreatic defect in cystic fibrosis. Acta Paediatr Scand Suppl 363, 41–44 (1989).

33 Fiorotto, R. & Strazzabosco, M. Pathophysiology of Cystic Fibrosis Liver Disease: A Channelopathy Leading to Alterations in Innate Immunity and in Microbiota. Cell Mol Gastroenterol Hepatol 8, 197–207 (2019).

34 Kim, S. Y. et al. Differentiating among conductive hearing loss conditions with wideband tympanometry. Auris Nasus Larynx 46, 43–49 (2019).

35 Merchant, G. R., Merchant, S. N., Rosowski, J. J. & Nakajima, H. H. Controlled exploration of the effects of conductive hearing loss on wideband acoustic immittance in human cadaveric preparations. Hear Res 341, 19–30 (2016).

36 Carraro, G. et al. Transcriptional analysis of cystic fibrosis airways at single-cell resolution reveals altered epithelial cell states and composition. Nat Med 27, 806–814 (2021).

37 Pruski, M. et al. Roles for IFT172 and Primary Cilia in Cell Migration, Cell Division, and Neocortex Development. Front Cell Dev Biol 7, 287 (2019).

38 Lee, M. S. et al. IFT46 plays an essential role in cilia development. Dev Biol 400, 248–257 (2015).

39 Di Gioia, S. A. et al. FAM161A, associated with retinitis pigmentosa, is a component of the cilia-basal body complex and interacts with proteins involved in ciliopathies. Hum Mol Genet 21, 5174–5184 (2012).

40 Bess, F. H., Dodd-Murphy, J. & Parker, R. A. Children with minimal sensorineural hearing loss: prevalence, educational performance, and functional status. Ear Hear 19, 339–354 (1998).

41 Yoshinaga-Itano, C. Benefits of early intervention for children with hearing loss. Otolaryngol Clin North Am 32, 1089–1102 (1999).

42 Fritze, W., Gotz, M., Stur, O. & Zweymuller, E. Hearing defects in cystic fibrosis. Z Kinderheilkd 114, 111–118 (1973).

43 Taylor, B., Evans, J. N. & Hope, G. A. Upper respiratory tract in cystic fibrosis. Ear-nose-throat survey of 50 children. Arch Dis Child 49, 133-136 (1974).

44 Berkhout, M. C. et al. Temporal bone pneumatization in cystic fibrosis: a correlation with genotype? Laryngoscope 124, 1682–1686 (2014).

45 Martins, L. M. et al. Hearing loss in cystic fibrosis. Int J Pediatr Otorhinolaryngol 74, 469–473 (2010).

46 Cepero, R. et al. Cystic fibrosis—an otolaryngologic perspective. Otolaryngology—Head and Neck Surgery 97, 356–360 (1987).

47 Boucher, R. C. Human airway ion transport. Part two. American Journal of Respiratory and Critical Care Medicine 150, 581–593 (1994).

48 Lang, F., Vallon, V., Knipper, M. & Wangemann, P. Functional significance of channels and transporters expressed in the inner ear and kidney. American Journal of Physiology-Cell Physiology 293, C1187–C1208 (2007).

49 Bazard, P. et al. Roles of Key Ion Channels and Transport Proteins in Age-Related Hearing Loss. International Journal of Molecular Sciences 22, 6158 (2021).

50 Parveen, S. et al. Evolution of Middle Ear Modelling Techniques: A Review. Cureus 13, e20829 (2021).

51 Voss, S. E., Nakajima, H. H., Huber, A. M. & Shera, C. A. in The Middle Ear Springer Handbook of Auditory Research Ch. Chapter 4, 67–91 (2013).

52 Thornton, J. L. et al. Conductive hearing loss induced by experimental middle-ear effusion in a chinchilla model reveals impaired tympanic membrane-coupled ossicular chain movement. J Assoc Res Otolaryngol 14, 451–464 (2013).

53 Keefe, D. H. & Simmons, J. L. Energy transmittance predicts conductive hearing loss in older children and adults. J Acoust Soc Am 114, 3217–3238 (2003).

54 Ryan, A. F. et al. Single-cell transcriptomes reveal a complex cellular landscape in the middle ear and differential capacities for acute response to infection. Frontiers in genetics 11, 358 (2020).

55 Burns, J. C. et al. Single-cell RNA-Seq resolves cellular complexity in sensory organs from the neonatal inner ear. Nat Commun 6, 8557 (2015).

56 MacArthur, C. J. et al. Otitis media impacts hundreds of mouse middle and inner ear genes. PloS one 8, e75213 (2013).

57 Hernandez, M. et al. The transcriptome of a complete episode of acute otitis media. BMC genomics 16, 1–16 (2015).

58 Gurr, A. et al. The ciliary beat frequency of middle ear mucosa in children with chronic secretory otitis media. Eur Arch Otorhinolaryngol 266, 1865–1870 (2009).

59 Arora, K. et al. Stabilizing rescued surface-localized deltaf508 CFTR by potentiation of its interaction with Na(+)/H(+) exchanger regulatory factor 1. Biochemistry 53, 4169–4179 (2014).

60 Penmatsa, H. et al. Compartmentalized cyclic adenosine 3’,5’-monophosphate at the plasma membrane clusters PDE3A and cystic fibrosis transmembrane conductance regulator into microdomains. Mol Biol Cell 21, 1097–1110 (2010).

61 Arora, K. et al. AC6 regulates the microtubule-depolymerizing kinesin KIF19A to control ciliary length in mammals. J Biol Chem 295, 14250–14259 (2020).

62 Francis, R. & Lo, C. Ex vivo method for high resolution imaging of cilia motility in rodent airway epithelia. J Vis Exp (2013).

63 Schneider, F., Draheirn, J., Kamberger, R. & Wallrabe, U. Process and material properties of polydimethylsiloxane (PDMS) for Optical MEMS. Sensor Actuat a-Phys 151, 95–99 (2009).

64 Muench, D. E. et al. Mouse models of neutropenia reveal progenitor-stage-specific defects. Nature 582, 109–114 (2020).

65 Venkatasubramanian, M. et al. Resolving single-cell heterogeneity from hundreds of thousands of cells through sequential hybrid clustering and NMF. Bioinformatics 36, 3773–3780 (2020).

66 DePasquale, E. A. K. et al. cellHarmony: cell-level matching and holistic comparison of single-cell transcriptomes. Nucleic Acids Res 47, e138 (2019).

67 Ransick, A. et al. Single-Cell Profiling Reveals Sex, Lineage, and Regional Diversity in the Mouse Kidney. Dev Cell 51, 399–413 e397 (2019).

